# PEFT-SP: Parameter-Efficient Fine-Tuning on Large Protein Language Models Improves Signal Peptide Prediction

**DOI:** 10.1101/2023.11.04.565642

**Authors:** Shuai Zeng, Duolin Wang, Dong Xu

## Abstract

Signal peptides (SP) play a crucial role in protein translocation in cells. The development of large protein language models (PLMs) provides a new opportunity for SP prediction, especially for the categories with limited annotated data. We present a Parameter-Efficient Fine-Tuning (PEFT) framework for SP prediction, PEFT-SP, to effectively utilize pre-trained PLMs. We implanted low-rank adaptation (LoRA) into ESM-2 models to better leverage the protein sequence evolutionary knowledge of PLMs. Experiments show that PEFT-SP using LoRA enhances state-of-the-art results, leading to a maximum MCC2 gain of 0.372 for SPs with small training samples and an overall MCC2 gain of 0.048. Furthermore, we also employed two other PEFT methods, i.e., Prompt Tunning and Adapter Tuning, into ESM-2 for SP prediction. More elaborate experiments show that PEFT-SP using Adapter Tuning can also improve the state-of-the-art results with up to 0.202 MCC2 gain for SPs with small training samples and an overall MCC2 gain of 0.030. LoRA requires fewer computing resources and less memory compared to Adapter, making it possible to adapt larger and more powerful protein models for SP prediction.

## 1 Introduction

Signal Peptides (SPs) are short amino acid sequences typically located in the N-terminals of nascent polypeptides and universally present in many proteins of a wide range of prokaryotic and eukaryotic organisms. Most SPs direct proteins to enter the secretory (Sec) pathway for translocation across the prokaryotic plasma membrane or the eukaryotic endoplasmic reticulum membrane. SPs containing a twin-arginine motif (R-R) target proteins to the twin-arginine translocation (Tat) pathway. The primary difference between the Sec and Tat pathways is that the Sec pathway transports proteins in unfolded conformation, whereas the Tat pathway translocates fully folded proteins.

Upon the successful translocation of the protein across the membrane, the SP is precisely cleaved at a specific cleavage site by signal peptidase (SPase). Subsequently, the mature protein is released on the trans side of the membrane [1]. The SPases are categorized into three groups: SPase I, II, and III (sometimes referred to as SPase IV [2]). SPase I cleaves general secretory signal peptides, whereas SPase II and SPase III are dedicated to cleavage signal peptides from lipoproteins and prepilin proteins, respectively. SPase I (Sec/SPI), SPase II (Sec/SPII), or SPase III (Sec/SPIII) can handle the processing of Sec substrates, whereas Tat substrates are exclusively processed by SPase I (Tat/SPI) or SPase II (Tat/SPII).

The signal peptidase cleavage site (CS) is recognized by SPase. Most SPs have a common tripartite structure, comprising a positively charged n-region, a central hydrophobic h-region spanning approximately 5-15 residues, and a c-region housing the CS for SPase I. Lipoprotein SPs, cleaved by SPase II, are recognized through the presence of a lipobox in the C-region. Prepilin SPs, subject to processing by SPase III, exclusively consist of a vital translocation-mediating region, as opposed to the conventional tripartite structure [3]. The amino acid composition and length of the SP regions exhibit diversity, to adapt to the specific requirements of various proteins within distinct cellular contexts. Although these SP regions are recognizable, the absence of clearly defined consensus motifs presents a significant challenge to SP prediction.

With the advances in machine learning and deep learning technologies, numerous applications for SP prediction have been developed and widely used in bioinformatics research. SignalP version 1-4 [4–7] are machine learning-based methods designed to predict Sec-translocated SPs cleaved by SPase I (Sec/SPI) and the corresponding CS locations. SPEPlip [8] employs a neural network approach combined with PROSITE patterns [9], allowing for the identification of SPs cleaved by SPase I (Sec/SPI) and lipoprotein SPs cleaved by SPase II (Sec/SPII). DeepSig [10] utilizes convolutional neural networks (CNN) and grammar-restrained conditional random fields to predict Sec-translocated SPs cleaved by SPase I and their cleavage sites. SignalP 5.0 [11] incorporates CNN and long short-term memory networks (LSTM) to predict Sec substrates cleaved by SPase I (Sec/SPI) or SPase II (Sec/SPII), as well as Tat substrates cleaved by SPase I (Tat/SPI). In contrast to its predecessors, SignalP 6.0 [12] stands out as a remarkable tool capable of predicting all five types of signal peptides (Sec/SPI, Sec/SPII, Tat/SPI, Tat/SPII, Sec/SPIII) through ProtTrans [13], a robust protein language model pre-trained on the UniRef100 dataset with a mask language model objective. Nevertheless, it is important to note that its performance in predicting SP with limited training samples leaves room for improvement.

Large protein language models (PLMs), such as ProTrans and ESM-1 [14], have become foundational tools for various biological modeling tasks related to proteins. Recently, ESM-2 increased the number of parameters in the Transformer model, which has led to substantial advancements in downstream protein prediction tasks[15]. The most common approach using pretrained PLMs for downstream tasks involves fine-tuning these models by updating all the parameters to leverage the information from the pre-trained model effectively.

While fine-tuning a model has proven to be a competitive strategy, the extensive fine-tuning process becomes impractical for large PLMs due to their significant computational and memory requirements. To tackle this challenge, a branch of research has emerged, focusing on Parameter-Efficient Fine-tuning (PEFT) for large language models, such as Adapter Tuning [16], Prompt Tuning [17], and Low-rank adaptation (LoRA) [18]. These techniques update introduced parameters within the pre-trained model, keeping all remaining parameters frozen during the training phase. The gradients of these frozen parameters are neither computed nor stored during back-propagation, substantially reducing computational and memory costs. Moreover, these approaches have demonstrated competitive performance compared to fine-tuning for various natural language processing (NLP) tasks.

In this paper, we present a novel SP prediction framework, PEFT-SP, designed to harness the capabilities of PLM for signal peptide and cleavage site prediction. PEFT-SP consists of the ESM-2 model, a linear Conditional Random Fields (CRF) model, and PEFT modules, including Adapter Tuning, Prompt Tuning, and Low-Rank adaptation (LoRA). The ESM-2 model serves as the backbone for encoding amino acid sequences and keeps frozen during the training phase. The CRF probabilistic model takes the representations generated by ESM-2 as input and predicts all five types of SPs and their corresponding CS. The PEFT method fine-tunes ESM-2 to suit the signal peptide prediction task better. Our framework is an end-to-end solution, focusing on optimizing parameters within CRF and PEFT modules exclusively. To demonstrate the effectiveness of our framework, we conducted a comprehensive performance comparison against existing SP predictors including the state-of-the-art tool, SignalP 6.0. Our results indicate that PEFT-SP using LoRA with ESM2-3B surpasses both the state-of-the-art tool and fine-tuned ESM-2 models across all five signal peptides. Notably, PEFT-SP using LoRA significantly improves performance in SP with limited training data. Additionally, we thoroughly investigated the performance of PEFP-SP using different PEFT methods with the ESM-2 model family for signal peptide prediction. In summary, our contributions to this study are as follows:

We introduced a novel signal peptide prediction framework, PEFT-SP, which leverages PEFT methods to finetune the ESM-2 model, resulting in enhanced predictions for both SP types and cleavage sites.

Our proposed method outperforms the current state-of-the-art model, SignalP 6.0, in two types of signal peptides with limited training samples, and it achieves comparable or superior performance in three other signal peptide types with larger training datasets.

We comprehensively evaluate ESM-2 fine-tuned and PEFT-SP using different combinations of the PEFT methods (including Prompt Tuning, Adapter Tuning, and LoRA) with the ESM-2 model family in the context of signal peptide prediction.

## 2 Methods

### 2.1 Pre-trained large protein language models

In recent years, several PLMs have emerged. For example, ProtTrans, ESM-1, and ESM-2 models are trained on sequences from the UniRef [19] protein sequence database using a masked language modeling objective. These models are specifically designed for protein feature extraction and can function as a foundation for fine-tuning in signal peptide prediction tasks. The state-of-the-art model SignalP 6.0 utilized ProtBert, derived from ProtTrans, as its backbone model. The ESM-2 model family, which encompasses varying model sizes ranging from 8 million parameters to a substantial 15 billion parameters, has shown remarkable performance in structure prediction. The ESM-2 model family, including ESM2-150M, ESM2-650M, and ESM2-3B, outperforms many protein language models from ProtTrans and the ESM-1 model family in protein sequence-related tasks. Additionally, we individually replaced the backbone in SignalP 6.0 with three backbones from the ESM-2 model family and fine-tuned them on the SP datasets. The ESM-2 model family also outperforms ProtBert (Supplementary Fig. 1) on the SP prediction. Hence, based on the solid performance in these protein sequence-related tasks, we employed the ESM-2 model family as the pre-trained backbone in our subsequent experiments. It is worth noting that while ESM-15B is available and exhibits excellent predictive capabilities, we have to exclude it from our study due to computational resource limitations.

**Fig. 1.**
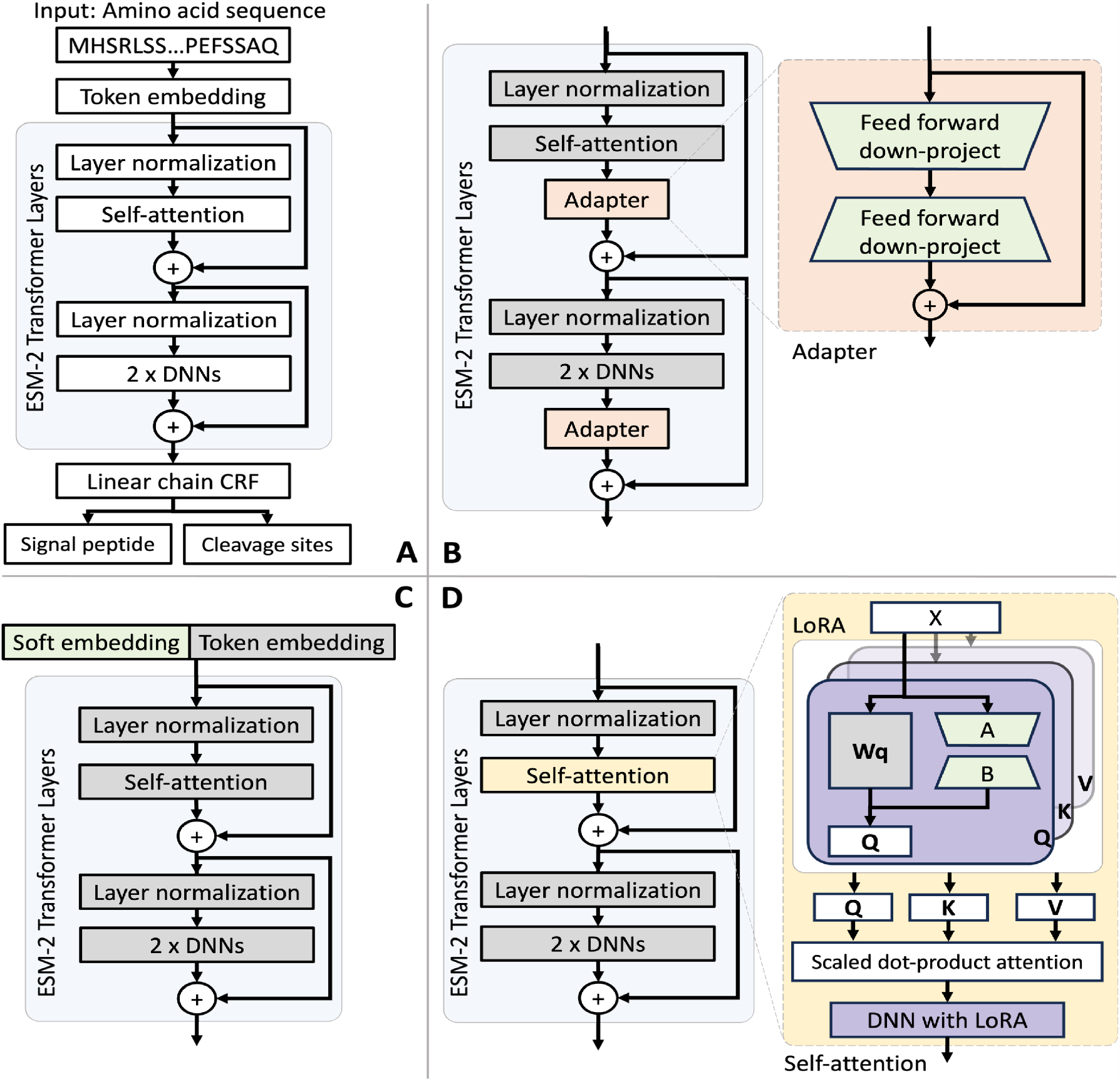
The architectures for the ESM-2 model and PEFT-SP using different PEFT modules. The light green modules are tunable during training, while the grey modules are fixed. (A) The ESM-2 backbone model uses amino acid sequences to SP and CS. (B) PEFT-SP using Adapter Tuning contains a bottleneck architecture. (C) PEFT-SP using Prompt Tuning appends soft embedding into token embedding. (D) PEFT-SP using LoRA adds trainable rank decomposition matrices into the self-attention layer.

Unlike existing signal peptide prediction models that require appending an organism identifier to the protein sequence, PEFT-SP with ESM-2 backbone (as shown in Fig. 1A) takes only the protein sequence *S* as input, encoding it into token embeddings. The token embeddings of the sequence are then fed into a stack of multiple Transformer layers, designed to learn contextual relationships between amino acids. Each Transformer layer consists of a self-attention mechanism and Position-wise Feed-Forward Networks (FFN) surrounded by separate residual connections. In the self-attention mechanism, the attention function processes the input feature *X* and transforms it into three different vectors of dimension *d*: query (*Q*), key (*K*), and value (*V*). This transformation uses three weight matrices: *W*_*q*_, *W*_*k*_, and *W*_*v*_. Subsequently, the Scaled Dot-Product Attention calculates the scaled attention score by performing a dot product between the *Q* and the *K* values associated with the amino acid token. It then converts this score into a probability distribution using the Softmax function:

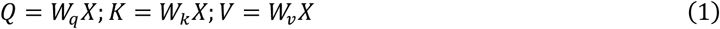

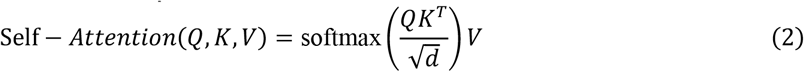

The output of the self-attention mechanism is a representation achieved through the weighted summation of values. The FFN module is constructed from two linear transformations, each activated by the Rectified Linear Unit (ReLU) function [20], yielding a sequence of hidden states. We remove the special tokens (CLS and SEP) introduced from the backbone and retain a sequence of hidden states *h* with the same length as the input sequence

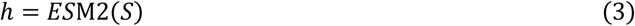

### 2.2 Parameter-Efficient Fine-tuning methods for ESM-2

Unlike the original configuration of Adapter Tuning and LoRA, which incorporate related modules into all Transformer layers, we specifically insert them into the bottommost *L* Transformer layers within the ESM-2 model. The idea comes from LLaMA-Adapter [21], which is designed to enhance the fine-tuning of representations with higher-level semantics.

Adapter Tuning. The Adapter Tuning [16] is the incorporation of adapter modules with a bottleneck architecture within the Transformer layer of the ESM-2 model. These adapter modules are introduced as distinct components, positioned after the projection phase following self-attention and after the two feed-forward layers. Each adapter module comprises a residual connection and a bottleneck architecture, which compress the input data into a bottleneck layer with reduced dimensionality and subsequently reconstructs the data to match the original input size. The fusion of the ESM-2 model with adapter modules is illustrated in Fig. 1B.

#### Prompt Tuning

The Prompt Tuning [17] method involves the addition of trainable embeddings, as “soft prompts,” into the sequence embeddings, which serve as inputs to the ESM-2 model. Considering the high sensitivity of prompt tuning to prompt initialization, prompts are initially set using embeddings of randomly selected amino acids. All parameters within the ESM-2 model remain fixed throughout the training process, while the soft prompts are continuously updated using gradients. Including soft prompts in the input sequence introduces extra hidden states generated by the ESM-2 model. To ensure that the length of the hidden states matches the sequence length, we omit the hidden states associated with the soft prompts. An overview of the ESM-2 model with prompt tuning is provided in Fig. 1C.

#### Low-Rank adaptation

LoRA [18] is built on the idea that trainable weights have a low “intrinsic rank”. This characteristic enables the weights to learn effectively, even when randomly projected into a smaller subspace. We perform lightweight fine-tuning of ESM-2 [15] by introducing trainable rank decomposition matrices into the Transformer architecture, implementing LoRA (as shown in Fig. 1D). Specifically, this reparameterization is applied to the projection matrices of the query, key, value, and FFN modules within the Transformer. The trained weight matrix, denoted as *W*_0_ ∈ *R*^*d*×*k*^, is coupled with a low-rank decomposition matrix Δ*W* = *BA*, where *B* ∈ *R*^*d*×r^, *A* ∈ *R*^*r*×*k*^, and *r* ≪ *min*(*d, k*). Both *W*_0_ and Δ*W* are simultaneously employed on the same input, and the output vectors they generate are combined elementwise. Hence, the functions for transforming X into a query (Q), key (K), and value (V) within the Transformer, as shown in Equation (1), undergo the following modifications:

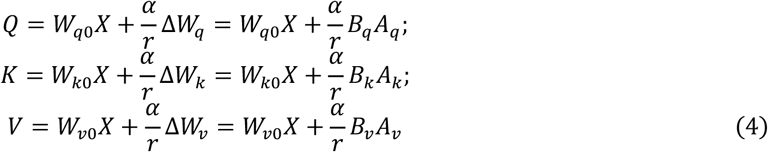

Where α is a scaling constant. We initialize *A* with random Gaussian initialization and *B* to zero, resulting in Δ*W* = *BA* to zero and preserving the original knowledge in the ESM-2 model at the beginning of training. During the training state, the pre-trained weight matrix *W*_0_ is kept frozen, and low-rank decomposition matrix Δ*W* is trainable by updating *A* and *B* with gradient. The hyperparameters *α* and *r* are determined through the hyperparameter tuning process.

### 2.3 Linear chain conditional random field

The linear chain CRF [22] is widely used in sequence labeling tasks, which can capture relationships between the labels in a sequence and the observed data. The linear chain CRF takes the sequence of hidden states *h* from the ESM-2 model and assigns regions of *y* for each sequence position based on the dependencies between neighboring states. The linear chain CRF can be written as:

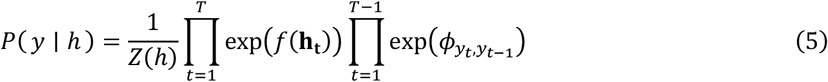

where *Z*(*h*) is a normalizing constant, *f*(·) denotes a linear transformation that prepares the emissions for the CRF, and *ϕ* represents the transition matrix governing the probability of *y*_*t*−1_ following *y*_*t*_. To accurately predict the defined regions of the SP class, which include the n-region, h-region, c-region, and twin-arginine motif, the *ϕ* transition matrix is constrained in the same way as SignalP 6.0 [12].

The Viterbi decoding process computes the most probable state sequence, encompassing the predicted SP class regions. For cleavage site prediction, the linear chain CRF predicts the regions of SP. The cleavage site (CS) is identified based on the last state of the region for the SP class within the most likely state sequence. The forward-backward algorithm calculates the marginal probabilities for each sequence position. To predict the type of signal peptide, the probability of a specific signal peptide type is calculated by summing the marginal probabilities associated with all states belonging to a particular type and then dividing by the sequence length:

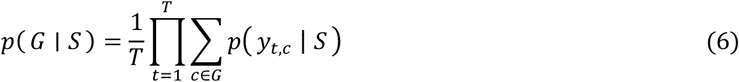

### 2.4 Loss function and regularization term

We constructed the loss function and regularization term following the approach of SignalP 6.0. We treated the SP region prediction task as a multilabel classification problem in the training stage. Specifically, positions around the boundaries of the regions were labeled as multilabel. For instance, a position near both the n-region and h-region was designated as both the n-region and h-region, reflecting the absence of a strict definition for region borders.

The loss function is derived from the negative log-likelihood of the linear chain CRF. At a specific position, the set of ground-truth labels is denoted as *M*_*t*_. The formulation of the loss function is as follows:

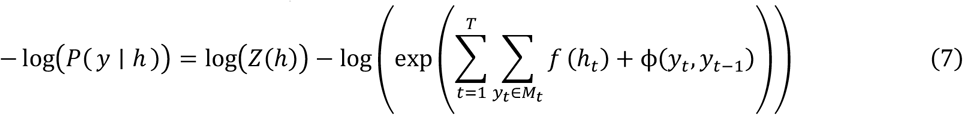

The regularization aims to promote diverse amino acid compositions within the three SP regions (n-region, h-region, c-region). We sum the marginal probabilities of all CRF states *c* belonging to region *r* for all positions with amino acid *a*, yielding a vector of *score*_*r*_ for each region *r*. Diversity scores are then calculated using cosine similarity for the vectors of scores between the n-region and h-region, as well as between the h-region and c-region, for each sequence. The mean of these diversity scores across all sequences is incorporated into the loss function with a scaling factor.

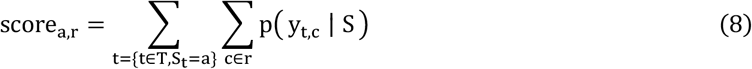

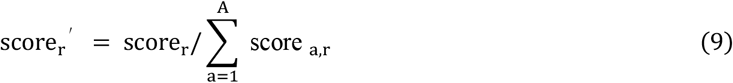

### 2.5 Model evaluation

We applied the same metrics used in baseline models, specifically the Matthews correlation coefficient (MCC), commonly employed in other signal peptide (SP) prediction methods for a fair comparison. Considering that most existing methods involve binary classification for distinguishing SP from non-SP, we calculated MCC metrics (MCC1) using a dataset where negative samples comprised transmembrane and soluble proteins. In addition, we computed MCC metrics (MCC2) on a dataset where a specific SP type was designated as the positive sample, with all other SP types and non-SP as the negative samples.

The precision and recall are used to assess the performance of CS prediction within a tolerance window of up to 3. Precision is the ratio of correct CSs to the total number of predicted CSs, while recall is the ratio of correctly predicted CSs to the total number of true CSs. For both metrics, a CS prediction was deemed accurate only if the predicted SP was also correct.

## 3 Experiment

### SP dataset

We utilized the benchmark dataset from SignalP 6.0, which includes a diverse set of protein sequences. The dataset consists of 3,352 Sec/SPI, 2,261 Sec/SPII, 113 Sec/SPIII, 595 Tat/SPI, 36 Tat/SPII, 16,421 intracellular sequences, and 2,615 transmembrane sequences. The SP types with limited training samples are Sec/SPIII and Tat/SPII. Each protein sequence in the dataset is accompanied by information about its SP type and region labels at each position, with the final label associated with the SP type indicating the CS. The dataset was initially acquired from four organism groups: Archaea, Eukarya, Gram-positive, and Gram-negative bacteria. The dataset was partitioned into three subsets to ensure fairness and robustness, with similar sequences grouped within each partition but no significant sequence similarity between proteins of different partitions. Furthermore, each partition was meticulously balanced across the four organism groups. Our experimental design involved a three-fold nested cross-validation process encompassing a two-fold inner loop and a three-fold outer loop. This configuration resulted in six distinct test sets. The distribution of the number of samples for SP type within each organism group is depicted in Supplementary Fig. 2.

**Fig. 2.**
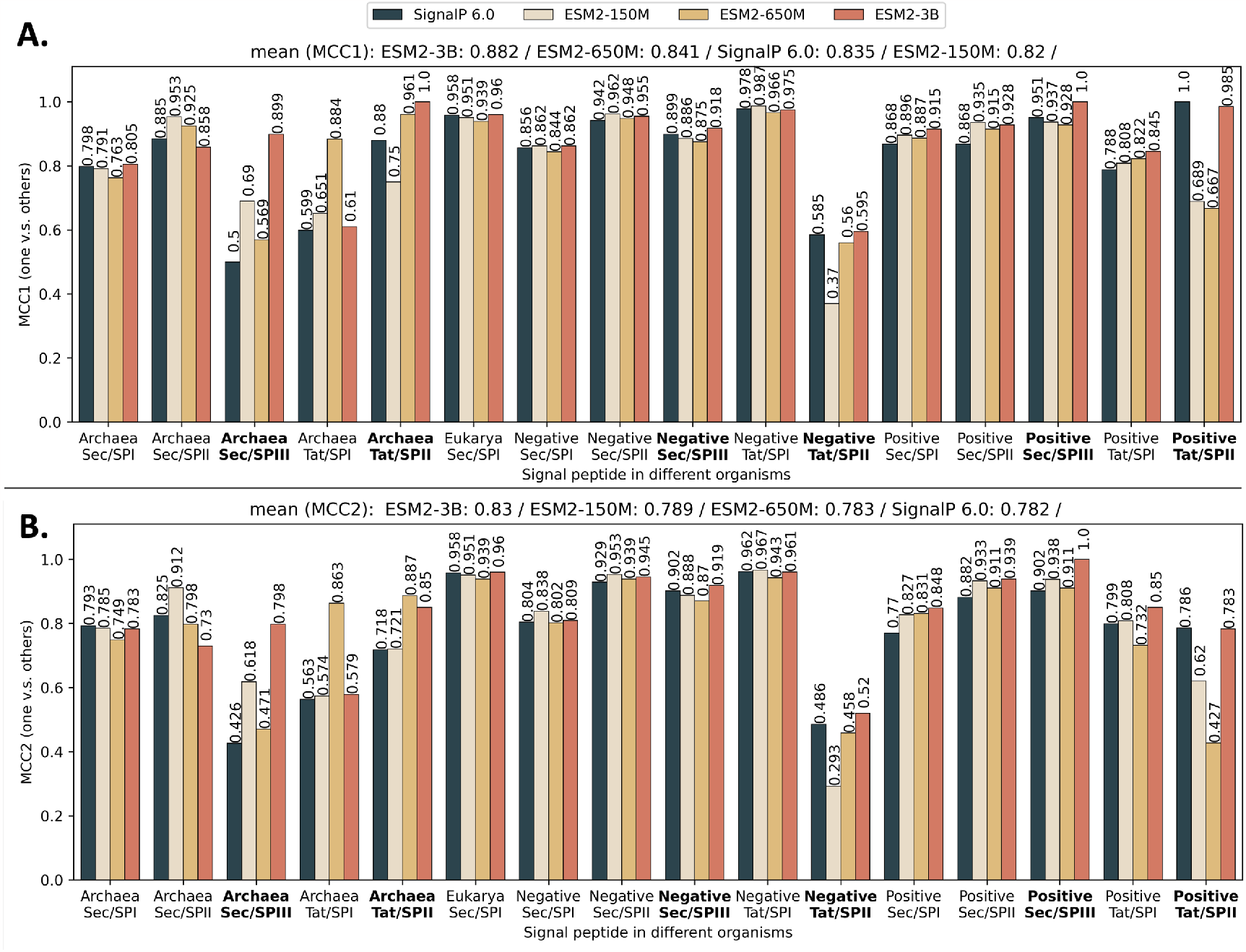
PEFT-SP using LoRA and SignalP 6.0 performance in terms of MCC score for each SP type across different organisms. The bold text in the x-axis represents the SP type with small training samples. The MCC1 and MCC2 scores are shown along with the bars. The sorted mean for MCC1 and MCC2 are listed at the top. (A) MCC1 scores performance on negative class composed of soluble and transmembrane proteins. (B) MCC2 scores performance on negative class comprising soluble and transmembrane proteins and other SP types.

### Experiment setting

We employed a pre-trained PLM from the ESM-2 [15] model family as the backbone for our model, comprising ESM2-150M, ESM2-650M, and ESM2-3B. Throughout the training process, the backbone model remains frozen. In contrast, the linear chain CRF and all parameters introduced by our PEFT methods, including LoRA [18], Prompt Tuning [17], and Adapter Tuning [16], were trainable. The model was trained end-to-end with the Adamax [23] optimizer. For model selection, we computed a combined MCC2 score for SP prediction and an MCC score for CS prediction. The best model was chosen from the validation sets based on the mean of these evaluation values. To optimize hyperparameters, we utilized Gaussian optimization provided by Optuna [24]. The specific hyperparameters and their corresponding search ranges are detailed in Supplementary Table 1. All runs were trained on a Nvidia A100 GPU with a batch size of 20.

**Table 1.**
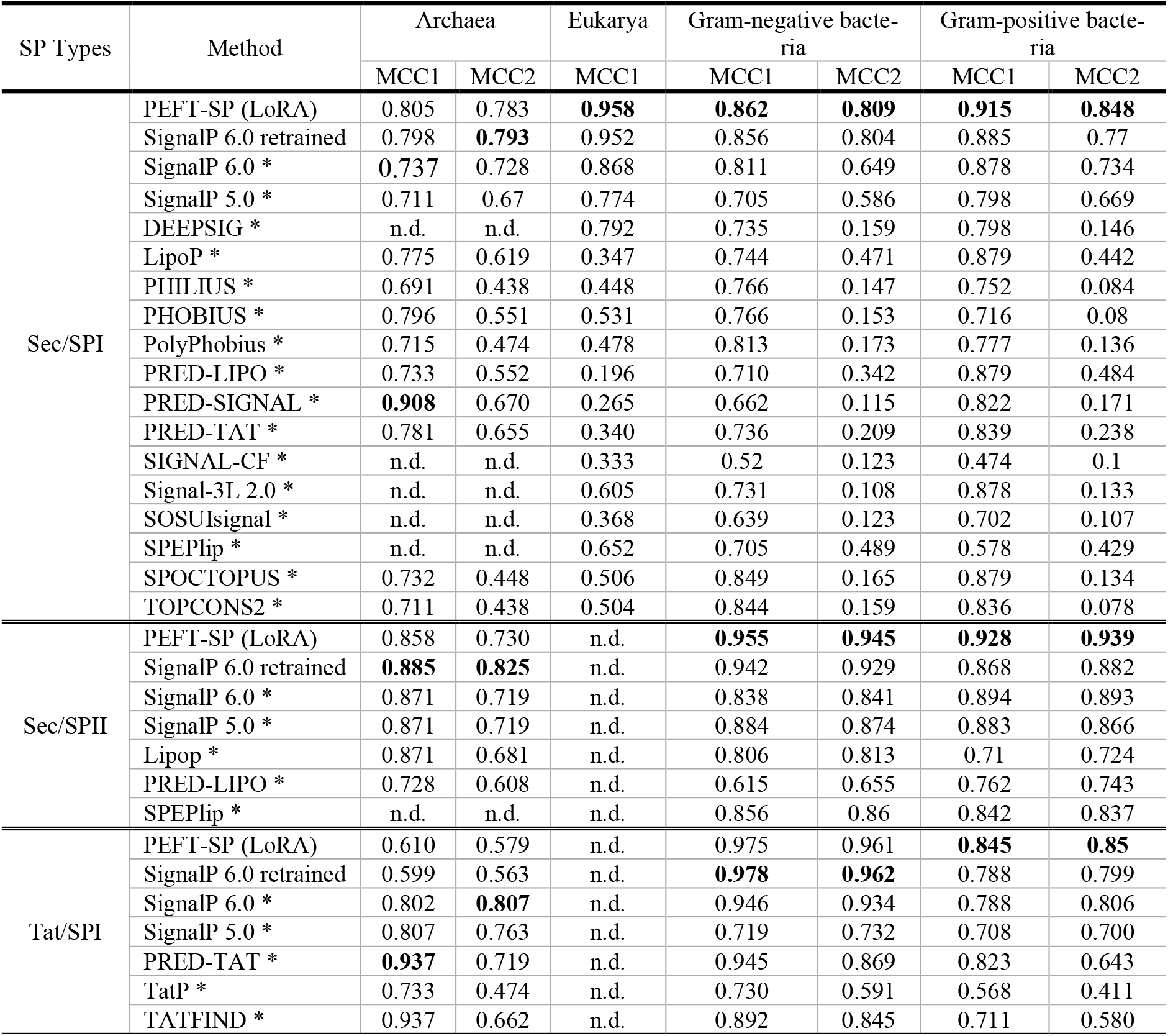
Benchmark results for SP prediction in Sec/SPI, Sec/SPII, and Tat/SPI. The values are the mean MCC1/MCC2 score across nest cross-validation. The bold value represents the highest MCC1/MCC2 score among the predictors in a particular SP type. The n.d. represents no data due to predictors not training on the data associated with SP type or the training data is available. The * denotes the performance reported in the SignalP 6.0.

We first compared our PEFT-SP using LoRA with the state-of-the-art model SignalP 6.0. Subsequently, we evaluated PEFT-SP using LoRA against all baseline models trained on the SP dataset with sufficient training samples. Finally, we extended our comparisons to fine-tuning and PEFT-SP using different combinations of PEFT methods (including Prompt Tuning, Adapter Tuning, and LoRA) with the ESM-2 model family. Our goal is to attain better performance in both SP and CS prediction compared to other existing methods.

## 4 Results

### 4.1 Comparisons with State-of-the-Art

As the well-trained models of SignalP 6.0 for nested cross-validation are not publicly available, we retrained it using the same datasets and default hyperparameters reported in the original paper. We employed PEFT-SP using LoRA for each model from the ESM-2 model family and trained them independently. We evaluated the MCC1 and MCC2 scores for each SP type within each organism group across test sets. Additionally, we calculated the mean MCC scores for MCC1 and MCC2 across all SP types and organisms.

PEFT-SP using LoRA with ESM2-3B backbone achieves the best performance (as shown in Fig.2). It consistently outperforms SignalP 6.0 in the SP types (Sec/SPIII and Tat/SPII) with limited training samples, except for Tat/SPII in Gram-positive bacteria. It achieves a maximum MCC1 gain of 0.399 and MCC2 gain of 0.372 in Sec/SPIII for Archaea. It attains a mean MCC1 improvement of 0.062 and a mean MCC2 improvement of 0.048. For SP types (Sec/SPI, Sec/SPII, and Tat/SPII) with sufficient training data, PEFT-SP using LoRA with ESM2-3B demonstrates superior or closely comparable performance to SignalP 6.0. We also visualized the confusion matrices for each organism group in Supplementary Fig.3. These matrices illustrate that PEFT-SP, using LoRA with the ESM2-3B backbone, exhibits strong performance in SP type prediction with fewer classification errors compared with SignalP 6.0.

We also computed precision and recall for CS prediction in PEFT-SP using LoRA and SignalP 6.0 (as shown in Fig. 3). Regarding precision, the PEFT-SP using LoRA with ESM2-3B backbone outperforms SignalP 6.0 in the Tat/SPII SP type, which is particularly notable given the limited training data for this type.

**Fig. 3.**
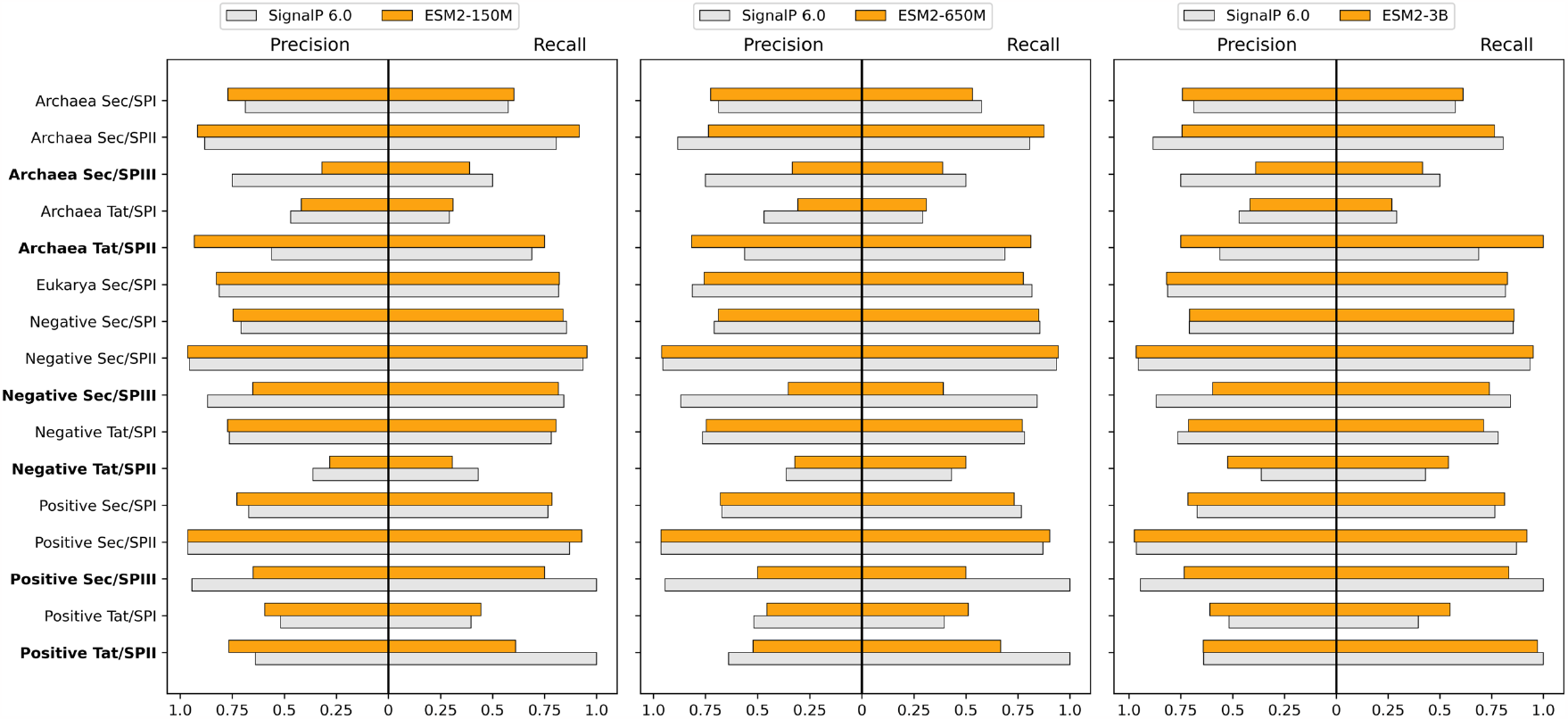
Results of PEFT-SP using LoRA and SignalP 6.0 in precision and recall for CS prediction across different organisms. The precision and recall are calculated within a tolerance window size of 0. The bold text in the x-axis represents the SP type with small training samples.

### 4.2 Comparisons with other baseline models

Considering the excellent performance of the PEFT-SP using LoRA with ESM2-3B [15] backbone, we compared it against all other baseline models. The performances for all baseline models were initially reported in SignalP 6.0 [12]. We included these performances in the benchmark. The benchmark also included the performance of SignalP 6.0, both when trained by our team and as reported in the original paper. The original baseline models were obtained from their publicly available web services, and all performance measurements were conducted on the same test sets generated through nested cross-validation. It is worth noting that, except for SignalP 6.0, the baseline models were trained on SP types with large training samples, and consequently, their performance regarding Sec/SPIII and Tat/SPII SP types has not been reported. Table 1 demonstrates that PEFT-SP using LoRA with ESM2-3B outperforms all baseline models in Sec/SPI for Eukarya, Sec/SPI and Sec/SPII for Gram-negative organisms, and all SP types for Gram-positive bacteria. Benchmark results for the recall of CS prediction in Sec/SPI, Sec/SPII and Tat/SPI at four tolerance windows can be found in Supplementary Tables 2-5.

**Table 2.** Benchmark results of MCC2 for SignalP 6.0, Fine-tuning ESM2-3B, and PEFT-SP using different PEFT methods with ESM2-3B backbone. The SP type indicated with the symbol † represents SP types with limited training samples. The bold value indicates the highest value for each SP type among all methods.

| Method / Backbone | - | Fine-tuning | Prompt Tuning | Adapter Tuning | LoRA |
| --- | --- | --- | --- | --- | --- |
| SP types | SignalP 6.0 | ESM2-3B | ESM2-3B | ESM2-3B | ESM2-3B |
| Archaea Sec/SPI | 0.793 | 0.771 | 0.777 | **0.825** | 0.783 |
| Archaea Sec/SPII | 0.825 | **0.864** | 0.509 | 0.783 | 0.730 |
| Archaea Sec/SPIII † | 0.426 | 0.724 | 0.500 | 0.351 | **0.798** |
| Archaea Tat/SPI | 0.563 | 0.564 | **0.653** | 0.538 | 0.579 |
| Archaea Tat/SPII † | 0.718 | 0.792 | 0.182 | 0.660 | **0.850** |
| Eukarya Sec/SPI | 0.958 | 0.948 | 0.954 | 0.954 | **0.960** |
| Negative Sec/SPI | 0.804 | **0.813** | 0.723 | 0.820 | 0.809 |
| Negative Sec/SPII | 0.929 | 0.946 | 0.886 | 0.950 | **0.945** |
| Negative Sec/SPIII † | 0.902 | **0.982** | 0.970 | 0.899 | 0.919 |
| Negative Tat/SPI | **0.962** | 0.902 | 0.853 | 0.899 | 0.961 |
| Negative Tat/SPII † | 0.486 | 0.358 | 0.325 | 0.405 | **0.520** |
| Positive Sec/SPI | 0.770 | 0.810 | 0.746 | 0.814 | **0.848** |
| Positive Sec/SPII | 0.882 | 0.908 | 0.833 | 0.911 | **0.939** |
| Positive Sec/SPIII † | 0.902 | **1.000** | 0.951 | 0.969 | **1.000** |
| Positive Tat/SPI | 0.799 | 0.746 | 0.590 | 0.752 | **0.850** |
| Positive Tat/SPII † | **0.786** | 0.603 | 0.148 | 0.669 | 0.783 |
| Mean (MCC2) | 0.781 | 0.796 | 0.663 | 0.762 | **0.830** |

### 4.3 Comparisons with fine-tuning and other PEFT methods

We compared PEFT-SP using different PEFT methods with ESM-3B, as well as SignalP 6.0 and finetuned ESM2-3B model. We trained all models independently with the same datasets generated from nest cross-validation. The performance of each model was measured using MCC2 across cross-validation.

Table 2 shows that the fine-tuning approach outperforms SignalP 6. This suggests that the ESM2-3B model holds promise as a potential candidate for other PEFT methods. The PEFT-SP using LoRA performs better than PEFT-SP using Prompt Tuning and Adapter Tuning regarding the mean MCC2. Moreover, the PEFT-SP using LoRA has less trainable parameters than fine-tuning and other PEFT methods, dramatically reducing the computing resource and memory storage. The number and percentage of trainable parameters for PEFT-SP are listed in Supplementary Table 6.

To comprehensively analyze the effectiveness of PEFT-SP using various PEFT methods with the ESM-2 model family, we benchmarked their results based on MCC1 and MCC2 for SP prediction (as presented in Supplementary Tables 7 and 8, respectively), and precision and recall for CS prediction (as presented in Supplementary Tables 9 and 10, respectively).

According to the benchmark results of MCC1 and MCC2, PEFT-SP using LoRA with ESM2-3B still performs best compared to other combinations. It is worth noting that PEFT-SP using Adapter Tuning with ESM-650M performs more promising than SignalP 6.0. It achieves a maximum MCC2 (MCC1) gain of 0.202 (0.250) in the SP types with limited training samples and a mean MCC2 (MCC1) gain of 0.03 (0.023) across all SP types.

## 5 Conclusion and Discussion

In this work, we presented PEFT-SP as a novel signal peptide prediction framework. It takes fragments of protein sequence as input without organism identifier. PEFT-SP using LoRA with ESM2-3B has demonstrated its capability to effectively handle SP types with limited training datasets and deliver performance comparable to or better than the baseline models across all SP types. The effectiveness of PEFT-SP using LoRA can be primarily attributed to the following factors: (1) PEFT-SP leverages the ESM2-3B backbone model, which captures the evolutionary aspects of protein sequences. (2) PEFT-SP employs LoRA, a lightweight fine-tuning method, to adapt PLM to SP prediction while preserving the high quality of the PLM.

We explored fine-tuning and different PEFT methods with the ESM2 family model for SP prediction. The finetuned models from the ESM2 model family perform better than the SignalP 6.0, suggesting ESM2 models are potential models for PEFT methods. Although Prompt Tuning has shown superior performance compared to the state-of-the-art in numerous tasks, we did not observe the same level of performance in SP prediction, possibly due to the ESM2-3B model not being sufficiently large. PEFT-SP using Adapter Tuning outperforms SignalP 6.0 but introduces a massive number of training parameters compared to PEFT-SP using LoRA.

To our best knowledge, this is the first study to explore the effectiveness of PLM using the PEFT approach for SP prediction tasks. There are many directions for future work: 1) PEFT-SP with combining PEFT methods, such as integrating PEFT-SP using LoRA and Adapter Tuning, potentially yields complementary improvements. 2) Modifying PEFT-SP to enhance interpretability, thereby unveiling the underlying motifs associated with SP. 3) Exploring the use of structure-aware PLM models as backbones, incorporating protein structure information to enhance SP prediction further. Our PEFT approach may be applicable to other protein predictions.

## Acknowledgments and Funding

We wish to thank Fei He and Yuexu Jiang for their useful discussions. This work was funded by the National Institutes of Health [R35-GM126985] and the National Science Foundation [DBI-2145226]. Funding for open access charge: National Science Foundation.

## Availability of Data and Materials

The SP dataset was downloaded from the web service of SignalP 6.0. The source code of PEFT-SP and trained models are publicly available at https://github.com/shuaizengMU/PEFT-SP.

## Notes

### Competing Interest Statement

The authors have declared no competing interest.

