## Supplementary Information for "PEFT-SP: Parameter-Efficient Fine-Tuning on Large Protein Language Models Improves Signal Peptide Prediction"


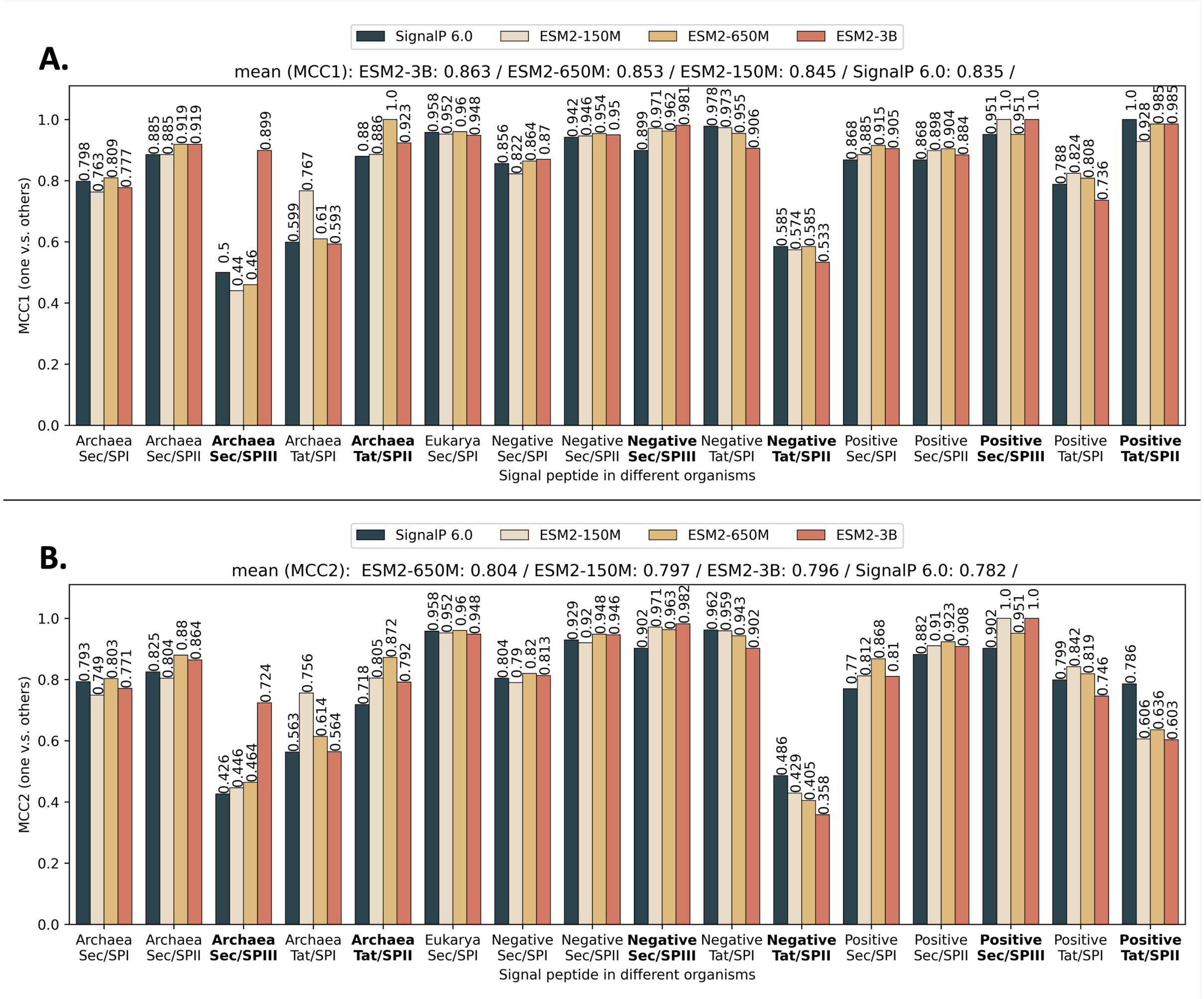


**Supplementary Fig. 1.** Results of finetuned SignalP 6.0 and finetuned ESM-2 model family with linear chain CRF in terms of MCC1 and MCC2 for SP prediction. The bold text in the x-axis represents the SP type with small training samples. The MCC1 and MCC2 scores are shown along with the bars. The sorted mean for MCC1 and MCC2 are listed at the top. (A) MCC1 scores performance on negative class composed of soluble and transmembrane proteins. (B) MCC2 scores performance on negative class comprising soluble and transmembrane proteins and other SP types.


* To whom correspondence should be addressed.

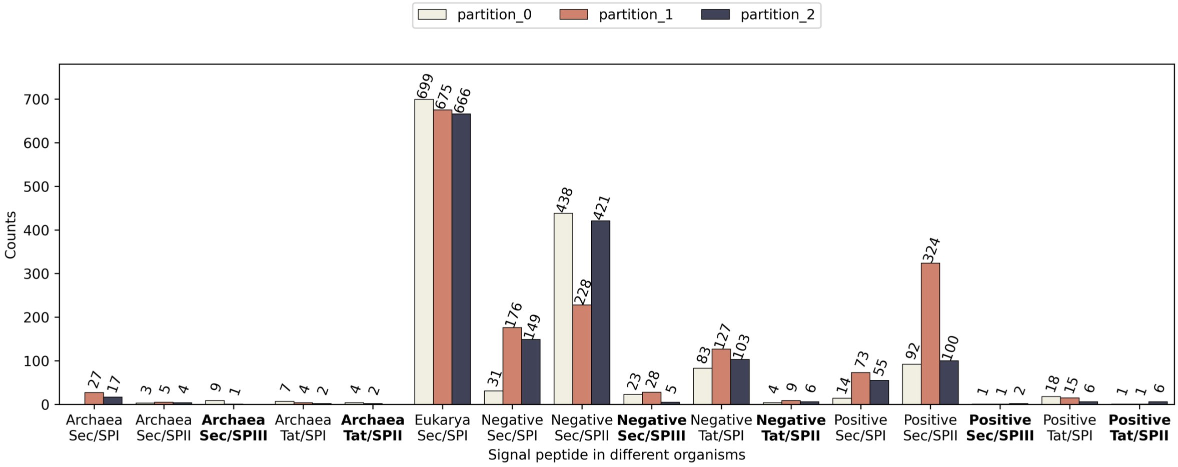


**Supplementary Fig. 2.** Distribution of the number of samples for SP type within each organism group. The partitions of the dataset are used for the nested cross-validation. The bold text in the x-axis represents the SP type with limited data.

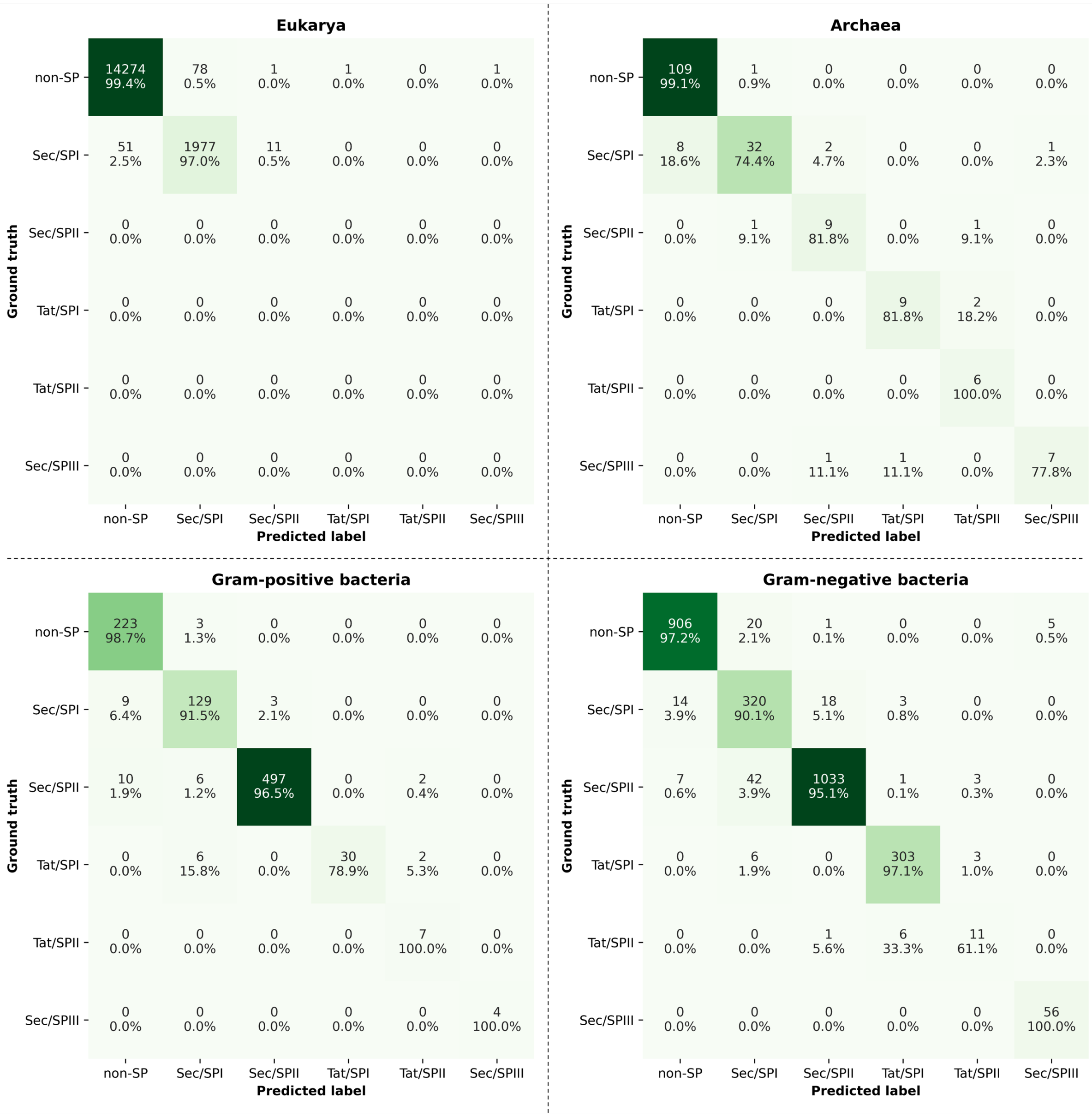


**Supplementary Fig. 3.** The cross-validated confusion matrices of PEFT-SP using LoRA with ESM2-3B across all organism groups. The non-SP group consists of sequences from both soluble and transmembrane proteins. Percentages are calculated row-wise to represent the proportion of correctly identified ground truth labels. The darker color represents a larger number of samples.

**Supplementary Table 1.** List of hyperparameters for each PEFT method, with the ranges specifying the search space for hyperparameter tuning.

| Methods | Hyperparameters | Ranges | Description |
|---|---|---|---|
| Adapter Tuning | num_last_layers | [1, number of Transformers in ESM-2 model] | Number of layers to add adapter modules at the end of ESM-2. |
| | learning_rate | [0.00001, 0.001] | Learning rate for Adapter and linear chain CRF. |
| Prompt Tuning | num_soft_prompt | [1, 70] | Number of soft prompts to add into the token embedding. |
| | learning_rate | [0.00001, 0.001] | Learning rate for soft prompt and linear chain CRF. |
| LoRA | r | [1,8] | Rank of decomposition matrix. |
| | α | [1,8] | Scaling const for decomposition matrix. |
| | num_last_layers | [1, number of Transformers in ESM-2 model] | Number of layers to add LoRA modules at the end of ESM-2. |
| | learning_rate | [0.00001, 0.001] | Learning rate for LoRA and linear chain CRF. |

**Supplementary Table 2.** Benchmark results for the recall of CS prediction in Sec/SPI at four tolerance windows. The PEFT-SP (LoRA) represents PEFT-SP using LoRA with ESM2-3B backbone. The bold value represents the highest recall among the predictors in a particular tolerance window. The n.d. represents no data due to predictors not training on the data associated with SP type or the training data is unavailable. The * denotes the performance reported in the SignalP 6.0.

| CS recall | Archaea | | | | Eukarya | | | | Gram - negative bacteria | | | | Gram - positive bacteria | | | |
|---|---|---|---|---|---|---|---|---|---|---|---|---|---|---|---|---|
| Method | ±0 | ±1 | ±2 | ±3 | ±0 | ±1 | ±2 | ±3 | ± 0 | ±1 | ± 2 | ±3 | ± 0 | ±1 | ±2 | ±3 |
| PEFT-SP (LoRA) | 0.612 | 0.679 | 0.679 | 0.708 | **0.827** | **0.866** | **0.901** | **0.920** | 0.711 | 0.764 | 0.878 | 0.904 | 0.813 | 0.832 | 0.852 | 0.860 |
| SignalP 6.0 retrained | 0.575 | 0.632 | 0.632 | 0.671 | 0.817 | 0.858 | **0.900** | **0.920** | **0.782** | **0.839** | **0.892** | **0.914** | 0.766 | 0.779 | 0.794 | 0.812 |
| SignalP 6.0 * | 0.500 | 0.556 | 0.556 | 0.583 | 0.747 | 0.774 | 0.808 | 0.829 | 0.639 | 0.672 | 0.689 | 0.721 | 0.800 | 0.800 | 0.800 | 0.800 |
| SignalP 5.0 * | 0.389 | 0.472 | 0.472 | 0.528 | 0.630 | 0.651 | 0.705 | 0.760 | 0.508 | 0.574 | 0.656 | 0.672 | 0.733 | 0.733 | 0.733 | 0.733 |
| DEEPSIG * | n.d. | n.d. | n.d. | n.d. | 0.603 | 0.630 | 0.658 | 0.699 | 0.508 | 0.574 | 0.574 | 0.574 | 0.733 | 0.733 | 0.800 | 0.800 |
| Lipop * | 0.389 | 0.528 | 0.556 | 0.639 | 0.288 | 0.329 | 0.370 | 0.404 | 0.656 | 0.705 | 0.721 | 0.721 | 0.467 | 0.467 | 0.533 | 0.533 |
| PHILIUS * | 0.500 | 0.611 | 0.611 | 0.611 | 0.596 | 0.658 | 0.712 | 0.760 | 0.623 | 0.672 | 0.721 | 0.754 | 0.467 | 0.467 | 0.467 | 0.467 |
| PHOBIUS * | 0.472 | 0.583 | 0.611 | 0.639 | 0.637 | 0.671 | 0.699 | 0.753 | 0.557 | 0.656 | 0.721 | 0.738 | 0.467 | 0.467 | 0.467 | 0.467 |
| PolyPhobius * | 0.528 | 0.667 | 0.667 | 0.667 | 0.623 | 0.678 | 0.733 | 0.801 | 0.557 | 0.672 | 0.754 | 0.754 | 0.667 | 0.667 | 0.733 | 0.733 |
| PRED-LIPO * | 0.472 | 0.556 | 0.611 | 0.639 | 0.068 | 0.082 | 0.130 | 0.158 | 0.410 | 0.475 | 0.508 | 0.525 | **0.867** | **0.867** | **0.867** | **0.867** |
| PRED-SIGNAL * | **0.861** | **0.917** | **0.917** | **0.917** | 0.226 | 0.267 | 0.301 | 0.329 | 0.426 | 0.492 | 0.607 | 0.639 | 0.800 | 0.800 | 0.800 | 0.800 |
| PRED-TAT * | 0.556 | 0.694 | 0.750 | 0.778 | 0.370 | 0.445 | 0.500 | 0.548 | 0.656 | 0.721 | 0.754 | 0.770 | **0.867** | **0.867** | **0.867** | **0.867** |
| Signal-3L 2.0 * | n.d. | n.d. | n.d.∂ | n.d. | 0.644 | 0.671 | 0.719 | 0.753 | 0.607 | 0.639 | 0.672 | 0.705 | 0.733 | 0.733 | 0.800 | 0.800 |
| Signal3Lv2 * | n.d. | n.d. | n.d. | n.d. | 0.664 | 0.685 | 0.726 | 0.753 | 0.541 | 0.607 | 0.623 | 0.639 | 0.800 | 0.800 | 0.800 | 0.800 |
| SOSUIsignal * | n.d. | n.d. | n.d. | n.d. | 0.151 | 0.308 | 0.459 | 0.568 | 0.246 | 0.377 | 0.557 | 0.623 | 0.200 | 0.267 | 0.267 | 0.467 |
| SPEPlip * | n.d. | n.d. | n.d. | n.d. | 0.685 | 0.712 | 0.747 | 0.781 | 0.574 | 0.656 | 0.705 | 0.721 | 0.600 | 0.600 | 0.667 | 0.667 |
| SPOCTOPUS * | 0.333 | 0.389 | 0.417 | 0.472 | 0.384 | 0.514 | 0.678 | 0.747 | 0.426 | 0.656 | 0.820 | 0.869 | 0.600 | 0.667 | 0.733 | 0.867 |
| TOPCONS2 * | 0.389 | 0.528 | 0.556 | 0.583 | 0.329 | 0.452 | 0.596 | 0.692 | 0.443 | 0.541 | 0.656 | 0.689 | 0.267 | 0.333 | 0.333 | 0.400 |

**Supplementary Table 3.** Benchmark results for the precision of CS prediction in Sec/SPI at four tolerance windows. The PEFT-SP (LoRA) represents PEFT-SP using LoRA with ESM2-3B backbone. The bold value represents the highest precision among the predictors in a particular tolerance window. The n.d. represents no data due to predictors not training on the data associated with SP type or the training data is unavailable. The * denotes the performance reported in the SignalP 6.0.

| CS precision | Archaea | | | | Eukarya | | | | Gram - negative bacteria | | | | Gram - positive bacteria | | | |
|---|---|---|---|---|---|---|---|---|---|---|---|---|---|---|---|---|
| Method | ±0 | ±1 | ±2 | ±3 | ± 0 | ±1 | ±2 | ±3 | ± 0 | ±1 | ± 2 | ±3 | ± 0 | ±1 | ±2 | ±3 |
| PEFT-SP (LoRA) | **0.742** | **0.829** | **0.829** | **0.856** | **0.818** | **0.857** | 0.891 | 0.910 | **0.707** | **0.736** | **0.743** | **0.750** | **0.715** | **0.734** | **0.755** | **0.763** |
| SignalP 6.0 retrained | 0.687 | 0.763 | 0.763 | 0.803 | 0.813 | 0.854 | **0.895** | **0.916** | **0.708** | **0.735** | **0.743** | **0.747** | 0.671 | 0.684 | 0.700 | 0.710 |
| SignalP 6.0 * | 0.643 | 0.714 | 0.714 | 0.750 | 0.661 | 0.685 | 0.715 | 0.733 | 0.534 | 0.562 | 0.575 | 0.603 | 0.632 | 0.632 | 0.632 | 0.632 |
| SignalP 5.0 * | 0.519 | 0.630 | 0.630 | 0.704 | 0.514 | 0.531 | 0.575 | 0.620 | 0.378 | 0.427 | 0.488 | 0.500 | 0.500 | 0.500 | 0.500 | 0.500 |
| DEEPSIG * | n.d. | n.d. | n.d. | n.d. | 0.587 | 0.613 | 0.640 | 0.680 | 0.134 | 0.151 | 0.151 | 0.151 | 0.089 | 0.089 | 0.098 | 0.098 |
| LipoP * | 0.359 | 0.487 | 0.513 | 0.590 | 0.141 | 0.162 | 0.182 | 0.199 | 0.339 | 0.364 | 0.373 | 0.373 | 0.152 | 0.152 | 0.174 | 0.174 |
| PHILIUS * | 0.353 | 0.431 | 0.431 | 0.431 | 0.168 | 0.186 | 0.201 | 0.215 | 0.110 | 0.118 | 0.127 | 0.133 | 0.051 | 0.051 | 0.051 | 0.051 |
| PHOBIUS * | 0.340 | 0.420 | 0.440 | 0.460 | 0.245 | 0.258 | 0.268 | 0.289 | 0.099 | 0.117 | 0.129 | 0.132 | 0.051 | 0.051 | 0.051 | 0.051 |
| PolyPhobius * | 0.352 | 0.444 | 0.444 | 0.444 | 0.181 | 0.197 | 0.213 | 0.233 | 0.098 | 0.118 | 0.133 | 0.133 | 0.069 | 0.069 | 0.076 | 0.076 |
| PRED - LIPO * | 0.386 | 0.455 | 0.500 | 0.523 | 0.052 | 0.062 | 0.098 | 0.119 | 0.203 | 0.236 | 0.252 | 0.260 | 0.325 | 0.325 | 0.325 | 0.325 |
| PRED - SIGNAL * | 0.508 | 0.541 | 0.541 | 0.541 | 0.073 | 0.086 | 0.097 | 0.106 | 0.085 | 0.098 | 0.121 | 0.128 | 0.083 | 0.083 | 0.083 | 0.083 |
| PRED - TAT * | 0.426 | 0.532 | 0.574 | 0.596 | 0.080 | 0.097 | 0.109 | 0.119 | 0.133 | 0.147 | 0.153 | 0.157 | 0.101 | 0.101 | 0.101 | 0.101 |
| Signal - 3L 2.0 * | n.d. | n.d. | n.d. | n.d. | 0.103 | 0.108 | 0.115 | 0.121 | 0.104 | 0.110 | 0.115 | 0.121 | 0.067 | 0.067 | 0.074 | 0.074 |
| Signal3Lv2 * | n.d. | n.d. | n.d. | n.d. | 0.357 | 0.368 | 0.390 | 0.404 | 0.116 | 0.130 | 0.134 | 0.137 | 0.093 | 0.093 | 0.093 | 0.093 |
| SOSUIsignal * | n.d. | n.d. | n.d. | n.d. | 0.032 | 0.066 | 0.098 | 0.122 | 0.042 | 0.065 | 0.096 | 0.107 | 0.021 | 0.028 | 0.028 | 0.049 |
| SPEPlip * | n.d. | n.d. | n.d. | n.d. | 0.362 | 0.377 | 0.395 | 0.413 | 0.278 | 0.317 | 0.341 | 0.349 | 0.257 | 0.257 | 0.286 | 0.286 |
| SPOCTOPUS * | 0.240 | 0.280 | 0.300 | 0.340 | 0.127 | 0.170 | 0.224 | 0.247 | 0.070 | 0.107 | 0.134 | 0.142 | 0.062 | 0.068 | 0.075 | 0.089 |
| TOPCONS2 * | 0.275 | 0.373 | 0.392 | 0.412 | 0.110 | 0.151 | 0.199 | 0.231 | 0.078 | 0.095 | 0.115 | 0.121 | 0.029 | 0.036 | 0.036 | 0.043 |

**Supplementary Table 4.** Benchmark results for the recall and precision of CS prediction in Sec/SPII at four tolerance windows. The PEFT-SP (LoRA) represents PEFT-SP using LoRA with ESM2-3B backbone. The bold value represents the highest recall or precision among the predictors in a particular tolerance window. The n.d. represents no data due to predictors not training on the data associated with SP type or the training data is unavailable. The * denotes the performance reported in the SignalP 6.0.

| | Archaea | | | | Gram - negative bacteria | | | | Gram - positive bacteria | | | |
|---|---|---|---|---|---|---|---|---|---|---|---|---|
| Method | ± 0 | ± 1 | ± 2 | ± 3 | ± 0 | ± 1 | ± 2 | ± 3 | ± 0 | ± 1 | ± 2 | ± 3 |
| CS recall | | | | | | | | | | | | |
| PEFT-SP (LoRA) | 0.764 | 0.764 | 0.764 | 0.764 | **0.858** | **0.895** | **0.900** | **0.908** | 0.813 | 0.832 | 0.852 | 0.860 |
| SignalP 6.0 retrained | **0.806** | **0.806** | **0.806** | **0.806** | 0.855 | 0.886 | 0.892 | 0.898 | 0.766 | 0.779 | 0.794 | 0.812 |
| SignalP 6.0 * | 0.778 | 0.778 | 0.778 | 0.778 | 0.852 | 0.852 | 0.856 | 0.864 | **0.875** | **0.883** | **0.883** | **0.883** |
| SignalP 5.0 * | 0.778 | 0.778 | 0.778 | 0.778 | 0.895 | 0.895 | 0.895 | 0.907 | 0.900 | 0.900 | 0.900 | 0.900 |
| LipoP * | 0.778 | 0.778 | 0.778 | 0.778 | 0.837 | 0.837 | 0.837 | 0.837 | 0.700 | 0.700 | 0.700 | 0.700 |
| PRED - LIPO * | 0.556 | 0.556 | 0.556 | 0.556 | 0.646 | 0.646 | 0.646 | 0.646 | 0.767 | 0.767 | 0.767 | 0.767 |
| SPEPlip * | n.d. | n.d. | n.d. | n.d. | 0.887 | 0.887 | 0.891 | 0.891 | 0.850 | 0.850 | 0.850 | 0.850 |
| CS precision | | | | | | | | | | | | |
| PEFT-SP (LoRA) | 0.744 | 0.744 | 0.744 | 0.744 | 0.707 | 0.736 | 0.743 | 0.750 | 0.715 | 0.734 | 0.755 | 0.763 |
| SignalP 6.0 retrained | **0.883** | **0.883** | **0.883** | **0.883** | 0.708 | 0.735 | 0.743 | 0.747 | 0.671 | 0.684 | 0.700 | 0.710 |
| SignalP 6.0 * | 0.583 | 0.583 | 0.583 | 0.583 | 0.913 | 0.913 | 0.917 | 0.925 | 0.929 | 0.938 | 0.938 | 0.938 |
| SignalP 5.0 * | 0.583 | 0.583 | 0.583 | 0.583 | 0.895 | 0.895 | 0.895 | 0.907 | 0.931 | 0.931 | 0.931 | 0.931 |
| LipoP * | 0.636 | 0.636 | 0.636 | 0.636 | 0.951 | 0.951 | 0.951 | 0.951 | 0.955 | 0.955 | 0.955 | 0.955 |
| PRED - LIPO * | 0.714 | 0.714 | 0.714 | 0.714 | **0.954** | **0.954** | 0.954 | 0.954 | 0.939 | 0.939 | 0.939 | 0.939 |
| SPEPlip * | n.d. | n.d. | n.d. | n.d. | **0.954** | **0.954** | **0.958** | **0.958** | **0.962** | **0.962** | **0.962** | **0.962** |

Supplementary **Table 5.** Benchmark results for the recall and precision of CS prediction in Tat/SPI at four tolerance windows. The PEFT-SP (LoRA) represents PEFT-SP using LoRA with ESM2-3B backbone. The bold value represents the highest recall or precision among the predictors in a particular tolerance window. The n.d. represents no data due to predictors not training on the data associated with SP type or the training data is unavailable. The * denotes the performance reported in the SignalP 6.0.

| | Archaea | | | | Gram - negative bacteria | | | | Gram - positive bacteria | | | |
|---|---|---|---|---|---|---|---|---|---|---|---|---|
| Method | ± 0 | ± 1 | ± 2 | ± 3 | ± 0 | ± 1 | ± 2 | ± 3 | ± 0 | ± 1 | ± 2 | ± 3 |
| CS recall | | | | | | | | | | | | |
| PEFT-SP (LoRA) | 0.268 | 0.292 | 0.339 | 0.405 | 0.711 | 0.764 | 0.878 | 0.904 | **0.550** | **0.550** | **0.765** | **0.765** |
| SignalP 6.0 retrained | 0.292 | 0.363 | 0.363 | 0.429 | **0.782** | **0.839** | **0.892** | **0.914** | 0.396 | 0.457 | 0.669 | 0.669 |
| SignalP 6.0 * | **0.333** | **0.444** | 0.444 | 0.444 | 0.706 | 0.765 | 0.784 | 0.804 | 0.556 | 0.556 | 0.667 | 0.667 |
| SignalP 5.0 * | 0.222 | **0.444** | 0.444 | 0.444 | 0.412 | 0.451 | 0.490 | 0.490 | 0.167 | 0.222 | 0.222 | 0.278 |
| PRED - TAT * | **0.333** | **0.444** | **0.667** | **0.667** | 0.725 | 0.765 | 0.784 | 0.824 | 0.611 | 0.611 | 0.667 | 0.722 |
| TatP * | 0.222 | 0.333 | 0.444 | 0.444 | 0.588 | 0.608 | 0.608 | 0.627 | 0.333 | 0.333 | 0.389 | 0.389 |
| TATFIND * | n.d. | n.d. | n.d. | n.d. | n.d. | n.d. | n.d. | n.d. | n.d. | n.d. | n.d. | n.d. |
| CS precision | | | | | | | | | | | | |
| PEFT-SP (LoRA) | 0.417 | 0.444 | 0.514 | 0.618 | 0.712 | 0.763 | **0.873** | **0.899** | 0.611 | 0.611 | 0.889 | 0.889 |
| SignalP 6.0 retrained | **0.469** | **0.562** | **0.562** | **0.656** | **0.764** | **0.820** | 0.872 | 0.895 | 0.518 | 0.595 | **0.921** | **0.921** |
| SignalP 6.0 * | 0.375 | 0.500 | 0.500 | 0.500 | 0.679 | 0.736 | 0.755 | 0.774 | **0.714** | **0.714** | 0.857 | 0.857 |
| SignalP 5.0 * | 0.182 | 0.364 | 0.364 | 0.364 | 0.488 | 0.535 | 0.581 | 0.581 | 0.273 | 0.364 | 0.364 | 0.455 |
| PRED - TAT * | 0.231 | 0.308 | 0.462 | 0.462 | 0.638 | 0.672 | 0.690 | 0.724 | 0.458 | 0.458 | 0.500 | 0.542 |
| TatP * | 0.133 | 0.200 | 0.267 | 0.267 | 0.326 | 0.337 | 0.337 | 0.348 | 0.167 | 0.167 | 0.194 | 0.194 |
| TATFIND * | n.d. | n.d. | n.d. | n.d. | n.d. | n.d. | n.d. | n.d. | n.d. | n.d. | n.d. | n.d. |

**Supplementary Table 6.** List of the number and percentage of trainable parameters for PEFT-SP using different combinations of the ESM-2 model family and PEFT method.

| Backbone | PEFT method | Total number of parameters | Number of trainable parameters | Percentage of training parameters |
|---|---|---:|---:|---:|
| ESM2-150M | Fine-tuning | 148,166,757 | 148,165,314 | 100.00% |
| | Adapter Tuning | 157,639,077 | 9,497,480 | 6.02% |
| | Prompt Tuning | 148,201,317 | 59,720 | 0.04% |
| | LoRA | 148,678,757 | 512,000 | 0.34% |
| ESM2-650M | Fine-tuning | 651,093,537 | 651,092,094 | 100.00% |
| | Adapter Tuning | 687,236,897 | 36,192,200 | 5.27% |
| | Prompt Tuning | 651,163,937 | 119,240 | 0.02% |
| | LoRA | 653,796,897 | 2,703,360 | 0.41% |
| ESM2-3B | Fine-tuning | 2,839,103,277 | 2,839,103,277 | 100.00% |
| | Adapter Tuning | 3,029,417,517 | 190,410,440 | 6.29% |
| | Prompt Tuning | 2,839,249,197 | 242,120 | 0.0085% |
| | LoRA | 2,844,837,677 | 5,734,400 | 0.20% |

**Supplementary Table 7.** Benchmark results of MCC1 for SignalP 6.0, Fine-tuning ESM-2 model family, and PEFT-SP using different PEFT methods with the ESM-2 model family. The SP type indicated with the symbol † represents SP types with limited training samples. The bold value indicates the highest value for each SP type among all methods.

| **Method** | **Baseline** | **Fine-tuning** | | | **Prompt Tuning** | | | **Adapter Tuning** | | | **LoRA** | | |
|---|---|---|---|---|---|---|---|---|---|---|---|---|---|
| **Backbone** | **SignalP 6.0** | **ESM2-150M** | **ESM2-650M** | **ESM2-3B** | **ESM2-150M** | **ESM2-650M** | **ESM2-3B** | **ESM2-150M** | **ESM2-650M** | **ESM2-3B** | **ESM2-150M** | **ESM2-650M** | **ESM2-3B** |
| **Archaea Sec/SPI** | 0.798 | 0.763 | 0.809 | 0.777 | 0.711 | 0.798 | 0.791 | **0.842** | 0.823 | 0.841 | 0.791 | 0.763 | 0.805 |
| **Archaea Sec/SPII** | 0.885 | 0.885 | 0.919 | 0.919 | 0.765 | 0.925 | 0.732 | 0.925 | 0.857 | 0.885 | **0.953** | 0.925 | 0.858 |
| **Archaea Sec/SPIII †** | 0.500 | 0.440 | 0.460 | **0.899** | 0.320 | 0.562 | 0.500 | 0.750 | 0.750 | 0.500 | 0.690 | 0.569 | **0.899** |
| **Archaea Tat/SPI** | 0.599 | 0.767 | 0.610 | 0.593 | 0.497 | 0.564 | 0.735 | 0.626 | 0.751 | 0.532 | 0.651 | **0.884** | 0.610 |
| **Archaea Tat/SPII †** | 0.880 | 0.886 | **1.000** | 0.923 | 0.367 | 0.461 | 0.250 | 0.669 | **1.000** | 0.867 | 0.750 | 0.961 | **1.000** |
| **Eukarya Sec/SPI** | 0.958 | 0.952 | 0.960 | 0.948 | 0.921 | 0.935 | 0.954 | 0.951 | 0.949 | 0.954 | 0.951 | 0.939 | **0.960** |
| **Negative Sec/SPI** | 0.856 | 0.822 | 0.864 | 0.870 | 0.773 | 0.791 | 0.819 | 0.875 | 0.869 | **0.877** | 0.862 | 0.844 | 0.862 |
| **Negative Sec/SPII** | 0.942 | 0.946 | 0.954 | 0.950 | 0.866 | 0.933 | 0.911 | 0.945 | 0.941 | 0.955 | **0.962** | 0.948 | 0.955 |
| **Negative Sec/SPIII †** | 0.899 | 0.971 | 0.962 | **0.981** | 0.925 | 0.977 | 0.969 | 0.883 | 0.964 | 0.897 | 0.886 | 0.875 | 0.918 |
| **Negative Tat/SPI** | 0.978 | 0.973 | 0.955 | 0.906 | 0.834 | 0.951 | 0.865 | 0.968 | 0.976 | 0.900 | **0.987** | 0.966 | 0.975 |
| **Negative Tat/SPII †** | 0.585 | 0.574 | 0.585 | 0.533 | 0.178 | 0.492 | 0.393 | 0.428 | 0.595 | **0.601** | 0.370 | 0.560 | 0.595 |
| **Positive Sec/SPI** | 0.868 | 0.885 | 0.915 | 0.905 | 0.847 | 0.800 | 0.867 | **0.920** | 0.897 | 0.918 | 0.896 | 0.887 | 0.915 |
| **Positive Sec/SPII** | 0.868 | 0.898 | 0.904 | 0.884 | 0.742 | 0.814 | 0.820 | 0.921 | 0.880 | 0.892 | **0.935** | 0.915 | 0.928 |
| **Positive Sec/SPIII †** | 0.951 | **1.000** | 0.951 | **1.000** | **1.000** | **1.000** | 0.951 | 0.928 | 0.969 | 0.969 | 0.937 | 0.928 | **1.000** |
| **Positive Tat/SPI** | 0.788 | 0.824 | 0.808 | 0.736 | 0.705 | 0.788 | 0.614 | 0.787 | 0.781 | 0.743 | 0.808 | 0.822 | **0.845** |
| **Positive Tat/SPII †** | **1.000** | 0.928 | 0.985 | 0.985 | 0.333 | 0.233 | 0.233 | 0.761 | 0.733 | **1.000** | 0.689 | 0.667 | 0.985 |
| **mean** | 0.835 | 0.845 | 0.852 | 0.863 | 0.674 | 0.752 | 0.713 | 0.824 | 0.858 | 0.833 | 0.820 | 0.841 | **0.882** |

**Supplementary Table 8.** Benchmark results of MCC2 for SignalP 6.0, Fine-tuning ESM-2 model family, and PEFT-SP using different PEFT methods with ESM-2 model family. The SP type indicated with the symbol † represents SP types with limited training samples. The bold value indicates the highest value for each SP type among all methods.

| **Method** | **Baseline** | **Fine-tuning** | | | **Prompt Tuning** | | | **Adapter Tuning** | | | **LoRA** | | |
|---|---|---|---|---|---|---|---|---|---|---|---|---|---|
| **Backbone** | **SignalP 6.0** | **ESM2-150M** | **ESM2-650M** | **ESM2-3B** | **ESM2-150M** | **ESM2-650M** | **ESM2-3B** | **ESM2-150M** | **ESM2-650M** | **ESM2-3B** | **ESM2-150M** | **ESM2-650M** | **ESM2-3B** |
| **Archaea Sec/SPI** | 0.793 | 0.749 | 0.803 | 0.771 | 0.689 | 0.784 | 0.777 | 0.820 | 0.802 | **0.825** | 0.785 | 0.749 | 0.783 |
| **Archaea Sec/SPII** | 0.825 | 0.804 | 0.880 | 0.864 | 0.541 | 0.723 | 0.509 | 0.882 | 0.798 | 0.783 | **0.912** | 0.798 | 0.730 |
| **Archaea Sec/SPIII †** | 0.426 | 0.446 | 0.464 | 0.724 | 0.318 | 0.568 | 0.500 | 0.672 | 0.639 | 0.351 | 0.618 | 0.471 | **0.798** |
| **Archaea Tat/SPI** | 0.563 | 0.756 | 0.614 | 0.564 | 0.389 | 0.499 | 0.653 | 0.583 | 0.755 | 0.538 | 0.574 | **0.863** | 0.579 |
| **Archaea Tat/SPII †** | 0.718 | 0.805 | 0.872 | 0.792 | 0.295 | 0.455 | 0.182 | 0.596 | **0.920** | 0.660 | 0.721 | 0.887 | 0.850 |
| **Eukarya Sec/SPI** | 0.958 | 0.952 | 0.960 | 0.948 | 0.921 | 0.935 | 0.954 | 0.951 | 0.949 | 0.954 | 0.951 | 0.939 | **0.960** |
| **Negative Sec/SPI** | 0.804 | 0.790 | 0.820 | 0.813 | 0.695 | 0.737 | 0.723 | 0.823 | 0.829 | 0.820 | **0.838** | 0.802 | 0.809 |
| **Negative Sec/SPII** | 0.929 | 0.920 | 0.948 | 0.946 | 0.854 | 0.898 | 0.886 | 0.939 | 0.932 | 0.950 | **0.953** | 0.939 | 0.945 |
| **Negative Sec/SPIII †** | 0.902 | 0.971 | 0.963 | **0.982** | 0.927 | 0.978 | 0.970 | 0.883 | 0.964 | 0.899 | 0.888 | 0.870 | 0.919 |
| **Negative Tat/SPI** | 0.962 | 0.959 | 0.943 | 0.902 | 0.747 | 0.934 | 0.853 | 0.955 | 0.964 | 0.899 | **0.967** | 0.943 | 0.961 |
| **Negative Tat/SPII †** | 0.486 | 0.429 | 0.405 | 0.358 | 0.084 | 0.449 | 0.325 | 0.309 | 0.481 | 0.405 | 0.293 | 0.458 | **0.520** |
| **Positive Sec/SPI** | 0.770 | 0.812 | **0.868** | 0.810 | 0.662 | 0.725 | 0.746 | 0.845 | 0.814 | 0.814 | 0.827 | 0.831 | 0.848 |
| **Positive Sec/SPII** | 0.882 | 0.910 | 0.923 | 0.908 | 0.767 | 0.835 | 0.833 | 0.928 | 0.892 | 0.911 | 0.933 | 0.911 | **0.939** |
| **Positive Sec/SPIII †** | 0.902 | **1.000** | 0.951 | **1.000** | **1.000** | **1.000** | 0.951 | 0.873 | 0.969 | 0.969 | 0.938 | 0.911 | **1.000** |
| **Positive Tat/SPI** | 0.799 | 0.842 | 0.819 | 0.746 | 0.575 | 0.644 | 0.590 | 0.799 | 0.721 | 0.752 | 0.808 | 0.732 | **0.850** |
| **Positive Tat/SPII †** | **0.786** | 0.606 | 0.636 | 0.603 | 0.240 | 0.233 | 0.148 | 0.587 | 0.540 | 0.669 | 0.620 | 0.427 | 0.783 |
| **mean** | 0.781 | 0.797 | 0.804 | 0.796 | 0.606 | 0.712 | 0.663 | 0.778 | 0.811 | 0.762 | 0.789 | 0.783 | **0.830** |

**Table 9.** Benchmark results for the precision of CS prediction at four different tolerance windows. The bold value represents the highest precision among the predictors in a particular tolerance window. The SP type indicated with the symbol † represents SP types with limited training samples.

| Method/backbone | | Baseline | Fine-tuning | | | Prompt Tuning | | | Adapter Tuning | | | LoRA | | |
|---|---|---|---|---|---|---|---|---|---|---|---|---|---|---|
| **Organism** | **window** | **SignalP 6.0** | **ESM2-150M** | **ESM2-650M** | **ESM2-3B** | **ESM2-150M** | **ESM2-650M** | **ESM2-3B** | **ESM2-150M** | **ESM2-650M** | **ESM2-3B** | **ESM2-150M** | **ESM2-650M** | **ESM2-3B** |
| **Archaea Sec/SPI** | **±0** | 0.687 | 0.708 | 0.750 | 0.750 | 0.596 | 0.692 | 0.664 | 0.678 | 0.697 | 0.696 | **0.771** | 0.725 | 0.742 |
| | **±1** | 0.763 | 0.826 | 0.817 | **0.861** | 0.691 | 0.782 | 0.762 | 0.770 | 0.773 | 0.762 | 0.814 | 0.831 | 0.829 |
| | **±2** | 0.763 | 0.843 | 0.817 | **0.861** | 0.691 | 0.782 | 0.762 | 0.796 | 0.773 | 0.775 | 0.830 | 0.831 | 0.829 |
| | **±3** | 0.803 | 0.869 | 0.843 | **0.889** | 0.719 | 0.841 | 0.788 | 0.822 | 0.798 | 0.828 | 0.873 | 0.857 | 0.856 |
| **Archaea Sec/SPII** | **±0** | 0.883 | 0.864 | **0.917** | 0.889 | 0.597 | 0.606 | 0.540 | **0.917** | 0.850 | 0.778 | **0.917** | 0.736 | 0.744 |
| | **±1** | 0.883 | 0.864 | **0.917** | 0.889 | 0.597 | 0.606 | 0.540 | **0.917** | 0.850 | 0.778 | **0.917** | 0.736 | 0.744 |
| | **±2** | 0.883 | 0.864 | **0.917** | 0.889 | 0.597 | 0.606 | 0.540 | **0.917** | 0.850 | 0.778 | **0.917** | 0.736 | 0.744 |
| | **±3** | 0.883 | 0.864 | **0.917** | 0.889 | 0.597 | 0.606 | 0.564 | **0.917** | 0.850 | 0.778 | **0.917** | 0.736 | 0.744 |
| **Archaea Sec/SPIII †** | **±0** | 0.750 | **1.000** | 0.857 | 0.875 | 0.500 | 0.625 | 0.417 | 0.431 | 0.583 | 0.417 | 0.319 | 0.333 | 0.389 |
| | **±1** | 0.750 | **1.000** | 0.857 | 0.875 | 0.500 | 0.625 | 0.417 | 0.542 | 0.583 | 0.417 | 0.403 | 0.396 | 0.444 |
| | **±2** | 0.750 | **1.000** | 0.857 | 0.875 | 0.500 | 0.625 | 0.417 | 0.625 | 0.583 | 0.417 | 0.403 | 0.396 | 0.583 |
| | **±3** | 0.750 | **1.000** | 0.857 | 0.875 | 0.500 | 0.625 | 0.417 | 0.625 | 0.583 | 0.417 | 0.403 | 0.396 | 0.583 |
| **Archaea Tat/SPI** | **±0** | 0.469 | 0.400 | 0.500 | 0.427 | 0.113 | 0.287 | 0.209 | 0.323 | **0.550** | 0.525 | 0.418 | 0.306 | 0.417 |
| | **±1** | 0.562 | 0.492 | 0.583 | 0.521 | 0.146 | 0.350 | 0.321 | 0.469 | **0.683** | 0.617 | 0.547 | 0.398 | 0.444 |
| | **±2** | 0.562 | 0.492 | 0.625 | 0.521 | 0.235 | 0.433 | 0.321 | 0.469 | **0.683** | 0.617 | 0.547 | 0.417 | 0.514 |
| | **±3** | 0.656 | 0.567 | 0.771 | 0.635 | 0.247 | 0.483 | 0.370 | 0.625 | **0.833** | 0.721 | 0.659 | 0.500 | 0.618 |
| **Archaea Tat/SPII †** | **±0** | 0.562 | 0.854 | 0.783 | 0.754 | 0.393 | 0.417 | 0.190 | 0.822 | 0.867 | 0.583 | **0.933** | 0.817 | 0.750 |
| | **±1** | 0.688 | 0.854 | 0.783 | 0.754 | 0.393 | 0.417 | 0.190 | 0.822 | 0.867 | 0.583 | **0.933** | 0.817 | 0.750 |
| | **±2** | 0.688 | 0.854 | 0.783 | 0.754 | 0.393 | 0.417 | 0.190 | 0.822 | 0.867 | 0.583 | **0.933** | 0.817 | 0.750 |
| | **±3** | 0.688 | 0.854 | 0.783 | 0.754 | 0.393 | 0.417 | 0.190 | 0.822 | 0.867 | 0.583 | **0.933** | 0.900 | 0.750 |
| **Eukarya Sec/SPI** | **±0** | 0.813 | 0.828 | 0.827 | 0.817 | 0.796 | 0.816 | **0.838** | 0.811 | 0.821 | 0.834 | 0.826 | 0.756 | 0.818 |
| | **±1** | 0.854 | 0.863 | 0.864 | 0.855 | 0.831 | 0.849 | **0.870** | 0.850 | 0.858 | 0.869 | 0.863 | 0.808 | 0.857 |
| | **±2** | 0.895 | 0.890 | 0.897 | 0.885 | 0.861 | 0.876 | 0.897 | 0.881 | 0.888 | **0.898** | 0.896 | 0.858 | 0.891 |
| | **±3** | **0.916** | 0.908 | 0.912 | 0.901 | 0.876 | 0.890 | 0.912 | 0.900 | 0.903 | 0.913 | 0.911 | 0.880 | 0.910 |
| **Negative Sec/SPI** | **±0** | 0.708 | 0.709 | 0.704 | 0.690 | 0.580 | 0.642 | 0.626 | 0.705 | 0.731 | 0.706 | **0.746** | 0.688 | 0.707 |
| | **±1** | 0.735 | 0.741 | 0.728 | 0.716 | 0.601 | 0.660 | 0.640 | 0.740 | 0.751 | 0.731 | **0.774** | 0.720 | 0.736 |
| | **±2** | 0.743 | 0.752 | 0.740 | 0.731 | 0.610 | 0.666 | 0.646 | 0.746 | 0.756 | 0.739 | **0.787** | 0.734 | 0.743 |
| | **±3** | 0.747 | 0.759 | 0.748 | 0.738 | 0.612 | 0.670 | 0.648 | 0.754 | 0.759 | 0.744 | **0.794** | 0.738 | 0.750 |
| **Negative Sec/SPII** | **±0** | 0.955 | 0.936 | **0.972** | 0.970 | 0.896 | 0.898 | 0.908 | 0.968 | 0.958 | 0.969 | 0.965 | 0.960 | 0.965 |
| | **±1** | 0.956 | 0.936 | **0.972** | 0.970 | 0.896 | 0.898 | 0.910 | 0.968 | 0.958 | 0.969 | 0.966 | 0.960 | 0.965 |
| | **±2** | 0.956 | 0.936 | **0.972** | 0.970 | 0.896 | 0.899 | 0.910 | 0.968 | 0.958 | 0.969 | 0.966 | 0.961 | 0.965 |
| | **±3** | 0.957 | 0.937 | **0.972** | 0.971 | 0.897 | 0.900 | 0.912 | 0.969 | 0.959 | 0.969 | 0.967 | 0.961 | 0.966 |
| **Negative Sec/SPIII †** | **±0** | **0.869** | 0.753 | 0.854 | 0.837 | 0.641 | 0.600 | 0.668 | 0.810 | 0.751 | 0.789 | 0.652 | 0.352 | 0.597 |
| | **±1** | **0.869** | 0.753 | 0.854 | 0.837 | 0.641 | 0.600 | 0.668 | 0.810 | 0.751 | 0.789 | 0.696 | 0.518 | 0.597 |

| Method/backbone | | Baseline | Fine-tuning | | | Prompt Tuning | | | Adapter Tuning | | | LoRA | | |
|---|---|---|---|---|---|---|---|---|---|---|---|---|---|---|
| **Organism** | **window** | **SignalP 6.0** | **ESM2-150M** | **ESM2-650M** | **ESM2-3B** | **ESM2-150M** | **ESM2-650M** | **ESM2-3B** | **ESM2-150M** | **ESM2-650M** | **ESM2-3B** | **ESM2-150M** | **ESM2-650M** | **ESM2-3B** |
| | **±2** | **0.869** | 0.753 | 0.854 | 0.861 | 0.641 | 0.600 | 0.668 | 0.810 | 0.751 | 0.789 | 0.696 | 0.577 | 0.597 |
| | **±3** | **0.869** | 0.753 | 0.854 | 0.861 | 0.641 | 0.600 | 0.668 | 0.810 | 0.751 | 0.789 | 0.696 | 0.666 | 0.758 |
| **Negative Tat/SPI** | **±0** | 0.764 | 0.765 | **0.793** | 0.785 | 0.647 | 0.717 | 0.683 | 0.779 | 0.764 | 0.789 | 0.773 | 0.746 | 0.712 |
| | **±1** | 0.820 | 0.806 | 0.837 | **0.844** | 0.696 | 0.760 | 0.750 | 0.817 | 0.816 | 0.839 | 0.818 | 0.785 | 0.763 |
| | **±2** | 0.872 | 0.856 | 0.891 | **0.895** | 0.727 | 0.806 | 0.786 | 0.867 | 0.859 | 0.887 | 0.868 | 0.853 | 0.873 |
| | **±3** | 0.895 | 0.880 | 0.903 | **0.910** | 0.745 | 0.817 | 0.800 | 0.897 | 0.881 | 0.900 | 0.880 | 0.871 | 0.899 |
| **Negative Tat/SPII †** | **±0** | 0.362 | 0.301 | 0.266 | 0.322 | 0.056 | 0.167 | 0.175 | 0.346 | 0.311 | 0.326 | 0.282 | 0.321 | **0.525** |
| | **±1** | 0.438 | 0.301 | 0.266 | 0.322 | 0.056 | 0.167 | 0.175 | 0.346 | 0.311 | 0.326 | 0.282 | 0.321 | **0.525** |
| | **±2** | 0.438 | 0.301 | 0.266 | 0.322 | 0.056 | 0.177 | 0.175 | 0.346 | 0.311 | 0.326 | 0.282 | 0.321 | **0.525** |
| | **±3** | 0.438 | 0.301 | 0.266 | 0.322 | 0.056 | 0.177 | 0.175 | 0.346 | 0.311 | 0.326 | 0.282 | 0.321 | **0.525** |
| **Positive Sec/SPI** | **±0** | 0.671 | 0.723 | **0.758** | 0.668 | 0.520 | 0.580 | 0.611 | 0.705 | 0.684 | 0.674 | 0.728 | 0.679 | 0.715 |
| | **±1** | 0.684 | 0.735 | **0.768** | 0.686 | 0.524 | 0.588 | 0.622 | 0.720 | 0.709 | 0.696 | 0.740 | 0.691 | 0.734 |
| | **±2** | 0.700 | 0.752 | **0.787** | 0.702 | 0.541 | 0.601 | 0.633 | 0.736 | 0.726 | 0.711 | 0.763 | 0.710 | 0.755 |
| | **±3** | 0.710 | 0.757 | **0.792** | 0.712 | 0.549 | 0.606 | 0.638 | 0.741 | 0.731 | 0.717 | 0.771 | 0.724 | 0.763 |
| **Positive Sec/SPII** | **±0** | 0.964 | 0.972 | 0.977 | **0.980** | 0.931 | 0.933 | 0.941 | 0.974 | 0.968 | 0.978 | 0.964 | 0.964 | 0.974 |
| | **±1** | 0.964 | 0.972 | 0.977 | **0.980** | 0.933 | 0.933 | 0.941 | 0.974 | 0.968 | 0.978 | 0.964 | 0.964 | 0.974 |
| | **±2** | 0.965 | 0.972 | 0.977 | **0.980** | 0.933 | 0.933 | 0.941 | 0.974 | 0.968 | 0.978 | 0.964 | 0.964 | 0.974 |
| | **±3** | 0.965 | 0.972 | 0.977 | **0.980** | 0.933 | 0.933 | 0.941 | 0.974 | 0.968 | 0.978 | 0.964 | 0.964 | 0.974 |
| **Positive Sec/SPIII †** | **±0** | 0.944 | 0.917 | 0.917 | **1.000** | 0.595 | 0.647 | 0.508 | 0.889 | **1.000** | 0.944 | 0.650 | 0.500 | 0.733 |
| | **±1** | 0.944 | 0.917 | 0.917 | **1.000** | 0.595 | 0.647 | 0.508 | 0.889 | **1.000** | 0.944 | 0.817 | 0.583 | 0.733 |
| | **±2** | 0.944 | 0.917 | 0.917 | **1.000** | 0.595 | 0.647 | 0.508 | 0.889 | **1.000** | 0.944 | 0.817 | 0.750 | 0.733 |
| | **±3** | 0.944 | 0.917 | 0.917 | **1.000** | 0.595 | 0.647 | 0.508 | 0.889 | **1.000** | 0.944 | 0.817 | 0.750 | 0.900 |
| **Positive Tat/SPI** | **±0** | 0.518 | **0.687** | 0.598 | 0.634 | 0.369 | 0.301 | 0.267 | 0.542 | 0.553 | 0.623 | 0.594 | 0.455 | 0.611 |
| | **±1** | 0.595 | **0.687** | 0.630 | 0.634 | 0.369 | 0.320 | 0.297 | 0.553 | 0.553 | 0.665 | 0.616 | 0.476 | 0.611 |
| | **±2** | 0.921 | **0.975** | 0.882 | 0.903 | 0.485 | 0.478 | 0.486 | 0.906 | 0.769 | 0.958 | 0.912 | 0.727 | 0.889 |
| | **±3** | 0.921 | **0.975** | 0.905 | 0.958 | 0.559 | 0.478 | 0.486 | 0.906 | 0.769 | 0.958 | 0.912 | 0.727 | 0.889 |
| **Positive Tat/SPII †** | **±0** | 0.639 | 0.428 | 0.467 | 0.436 | 0.329 | 0.137 | 0.117 | 0.621 | 0.573 | 0.550 | **0.767** | 0.521 | 0.641 |
| | **±1** | 0.639 | 0.461 | 0.467 | 0.436 | 0.329 | 0.137 | 0.117 | 0.621 | 0.573 | 0.550 | **0.767** | 0.521 | 0.641 |
| | **±2** | 0.639 | 0.461 | 0.467 | 0.436 | 0.329 | 0.137 | 0.117 | 0.621 | 0.573 | 0.550 | **0.767** | 0.521 | 0.641 |
| | **±3** | 0.639 | 0.461 | 0.467 | 0.436 | 0.329 | 0.137 | 0.117 | 0.621 | 0.573 | 0.550 | **0.767** | 0.521 | 0.641 |
| **mean** | | 0.767 | 0.773 | **0.777** | 0.772 | 0.558 | 0.592 | 0.550 | 0.752 | 0.758 | 0.730 | 0.752 | 0.673 | 0.733 |

**Table 10.** Benchmark results for the recall of CS prediction at four different tolerance windows. The bold value represents the highest recall among the predictors in a particular tolerance window. The SP type indicated with the symbol † represents SP types with limited training samples.

| Method/backbone | | Baseline | Fine-tuning | | | Prompt Tuning | | | Adapter Tuning | | | LoRA | | |
|---|---|---|---|---|---|---|---|---|---|---|---|---|---|---|
| **Organism** | **window** | **SignalP 6.0** | **ESM2-150M** | **ESM2-650M** | **ESM2-3B** | **ESM2-150M** | **ESM2-650M** | **ESM2-3B** | **ESM2-150M** | **ESM2-650M** | **ESM2-3B** | **ESM2-150M** | **ESM2-650M** | **ESM2-3B** |
| **Archaea Sec/SPI** | **±0** | 0.575 | 0.547 | **0.640** | 0.570 | 0.490 | 0.575 | 0.575 | 0.612 | 0.625 | 0.611 | 0.603 | 0.531 | 0.612 |
| | **±1** | 0.632 | 0.623 | **0.697** | 0.651 | 0.557 | 0.642 | 0.651 | 0.688 | **0.697** | 0.673 | 0.642 | 0.618 | 0.679 |
| | **±2** | 0.632 | 0.632 | 0.697 | 0.651 | 0.557 | 0.642 | 0.651 | **0.706** | 0.697 | 0.682 | 0.651 | 0.618 | 0.679 |
| | **±3** | 0.671 | 0.662 | 0.727 | 0.680 | 0.586 | 0.690 | 0.680 | **0.736** | 0.727 | **0.736** | 0.690 | 0.647 | 0.708 |
| **Archaea Sec/SPII** | **±0** | 0.806 | 0.806 | 0.861 | 0.861 | 0.706 | 0.875 | 0.667 | 0.875 | 0.778 | 0.806 | **0.917** | 0.875 | 0.764 |
| | **±1** | 0.806 | 0.806 | 0.861 | 0.861 | 0.706 | 0.875 | 0.667 | 0.875 | 0.778 | 0.806 | **0.917** | 0.875 | 0.764 |
| | **±2** | 0.806 | 0.806 | 0.861 | 0.861 | 0.706 | 0.875 | 0.667 | 0.875 | 0.778 | 0.806 | **0.917** | 0.875 | 0.764 |
| | **±3** | 0.806 | 0.806 | 0.861 | 0.861 | 0.706 | 0.875 | 0.708 | 0.875 | 0.778 | 0.806 | **0.917** | 0.875 | 0.764 |
| **Archaea Sec/SPIII** | **±0** | 0.500 | 0.417 | 0.389 | **0.861** | 0.278 | 0.528 | 0.500 | 0.556 | 0.750 | 0.500 | 0.389 | 0.389 | 0.417 |
| | **±1** | 0.500 | 0.417 | 0.389 | **0.861** | 0.278 | 0.528 | 0.500 | 0.667 | 0.750 | 0.500 | 0.444 | 0.417 | 0.472 |
| | **±2** | 0.500 | 0.417 | 0.389 | **0.861** | 0.278 | 0.528 | 0.500 | 0.750 | 0.750 | 0.500 | 0.444 | 0.417 | 0.611 |
| | **±3** | 0.500 | 0.417 | 0.389 | **0.861** | 0.278 | 0.528 | 0.500 | 0.750 | 0.750 | 0.500 | 0.444 | 0.417 | 0.611 |
| **Archaea Tat/SPI** | **±0** | 0.292 | 0.333 | 0.292 | 0.226 | 0.155 | 0.226 | 0.286 | 0.220 | **0.351** | 0.268 | 0.310 | 0.310 | 0.268 |
| | **±1** | 0.363 | 0.405 | 0.339 | 0.274 | 0.179 | 0.274 | 0.405 | 0.315 | **0.446** | 0.315 | 0.405 | 0.405 | 0.292 |
| | **±2** | 0.363 | 0.405 | 0.363 | 0.274 | 0.250 | 0.345 | 0.405 | 0.315 | **0.446** | 0.315 | 0.405 | 0.429 | 0.339 |
| | **±3** | 0.429 | 0.470 | 0.446 | 0.339 | 0.292 | 0.387 | 0.488 | 0.423 | **0.571** | 0.381 | 0.488 | 0.512 | 0.405 |
| **Archaea Tat/SPII** | **±0** | 0.688 | 0.812 | **1.000** | 0.875 | 0.312 | 0.438 | 0.250 | 0.625 | **1.000** | 0.812 | 0.750 | 0.812 | **1.000** |
| | **±1** | 0.812 | 0.812 | **1.000** | 0.875 | 0.312 | 0.438 | 0.250 | 0.625 | **1.000** | 0.812 | 0.750 | 0.812 | **1.000** |
| | **±2** | 0.812 | 0.812 | **1.000** | 0.875 | 0.312 | 0.438 | 0.250 | 0.625 | **1.000** | 0.812 | 0.750 | 0.812 | **1.000** |
| | **±3** | 0.812 | 0.812 | **1.000** | 0.875 | 0.312 | 0.438 | 0.250 | 0.625 | **1.000** | 0.812 | 0.750 | 0.875 | **1.000** |
| **Eukarya Sec/SPI** | **±0** | 0.817 | 0.825 | 0.839 | 0.835 | 0.825 | 0.828 | 0.835 | 0.826 | 0.827 | **0.846** | 0.821 | 0.776 | 0.827 |
| | **±1** | 0.858 | 0.860 | 0.877 | 0.873 | 0.861 | 0.861 | 0.867 | 0.866 | 0.864 | **0.882** | 0.858 | 0.830 | 0.866 |
| | **±2** | 0.900 | 0.887 | 0.910 | 0.904 | 0.891 | 0.888 | 0.894 | 0.897 | 0.894 | **0.910** | 0.889 | 0.881 | 0.901 |
| | **±3** | 0.920 | 0.904 | 0.925 | 0.920 | 0.907 | 0.902 | 0.908 | 0.917 | 0.909 | **0.926** | 0.905 | 0.904 | 0.920 |
| **Negative Sec/SPI** | **±0** | 0.855 | 0.789 | 0.847 | 0.873 | 0.790 | 0.774 | 0.819 | 0.862 | 0.889 | **0.894** | 0.839 | 0.850 | 0.858 |
| | **±1** | 0.886 | 0.830 | 0.875 | 0.906 | 0.819 | 0.793 | 0.840 | 0.904 | 0.912 | **0.925** | 0.872 | 0.889 | 0.895 |
| | **±2** | 0.892 | 0.841 | 0.887 | 0.921 | 0.826 | 0.798 | 0.847 | 0.911 | 0.915 | **0.933** | 0.885 | 0.904 | 0.900 |
| | **±3** | 0.898 | 0.849 | 0.895 | 0.928 | 0.828 | 0.802 | 0.850 | 0.921 | 0.920 | **0.938** | 0.892 | 0.909 | 0.908 |
| **Negative Sec/SPII** | **±0** | 0.935 | 0.947 | 0.948 | 0.947 | 0.885 | 0.945 | 0.925 | 0.938 | 0.949 | 0.947 | **0.954** | 0.944 | 0.951 |
| | **±1** | 0.936 | 0.947 | 0.948 | 0.947 | 0.885 | 0.945 | 0.927 | 0.938 | 0.949 | 0.947 | **0.955** | 0.944 | 0.951 |
| | **±2** | 0.936 | 0.947 | 0.948 | 0.947 | 0.885 | 0.945 | 0.927 | 0.938 | 0.949 | 0.947 | **0.955** | 0.945 | 0.951 |
| | **±3** | 0.937 | 0.947 | 0.949 | 0.948 | 0.885 | 0.946 | 0.929 | 0.939 | 0.950 | 0.948 | **0.956** | 0.945 | 0.952 |
| **Negative Sec/SPIII** | **±0** | 0.842 | 0.993 | 0.952 | 0.959 | 0.928 | 0.978 | 0.964 | **1.000** | **1.000** | 0.899 | 0.816 | 0.391 | 0.739 |
| | **±1** | 0.842 | 0.993 | 0.952 | 0.959 | 0.928 | 0.978 | 0.964 | **1.000** | **1.000** | 0.899 | 0.859 | 0.591 | 0.739 |

| Method/backbone | | Baseline | Fine-tuning | | | Prompt Tuning | | | Adapter Tuning | | | LoRA | | |
|---|---|---|---|---|---|---|---|---|---|---|---|---|---|---|
| Organism | window | SignalP 6.0 | ESM2-150M | ESM2-650M | ESM2-3B | ESM2-150M | ESM2-650M | ESM2-3B | ESM2-150M | ESM2-650M | ESM2-3B | ESM2-150M | ESM2-650M | ESM2-3B |
| | ±2 | 0.842 | 0.993 | 0.952 | 0.993 | 0.928 | 0.978 | 0.964 | **1.000** | **1.000** | 0.899 | 0.859 | 0.765 | 0.739 |
| | ±3 | 0.842 | 0.993 | 0.952 | 0.993 | 0.928 | 0.978 | 0.964 | **1.000** | **1.000** | 0.899 | 0.859 | 0.859 | 0.900 |
| **Negative Tat/SPI** | ±0 | 0.782 | 0.788 | 0.781 | 0.716 | 0.656 | 0.772 | 0.649 | 0.776 | 0.792 | 0.712 | **0.805** | 0.771 | 0.711 |
| | ±1 | 0.839 | 0.833 | 0.828 | 0.773 | 0.698 | 0.817 | 0.703 | 0.816 | 0.847 | 0.763 | **0.852** | 0.815 | 0.764 |
| | ±2 | 0.892 | 0.882 | 0.879 | 0.816 | 0.732 | 0.867 | 0.738 | 0.865 | 0.891 | 0.802 | **0.904** | 0.886 | 0.878 |
| | ±3 | 0.914 | 0.906 | 0.891 | 0.828 | 0.754 | 0.878 | 0.752 | 0.894 | 0.913 | 0.815 | **0.916** | 0.902 | 0.904 |
| **Negative Tat/SPII** | ±0 | 0.431 | 0.505 | 0.523 | 0.458 | 0.097 | 0.347 | 0.375 | 0.375 | **0.542** | 0.486 | 0.306 | 0.500 | **0.542** |
| | ±1 | 0.523 | 0.505 | 0.523 | 0.458 | 0.097 | 0.347 | 0.375 | 0.375 | **0.542** | 0.486 | 0.306 | 0.500 | **0.542** |
| | ±2 | 0.523 | 0.505 | 0.523 | 0.458 | 0.097 | 0.384 | 0.375 | 0.375 | **0.542** | 0.486 | 0.306 | 0.500 | **0.542** |
| | ±3 | 0.523 | 0.505 | 0.523 | 0.458 | 0.097 | 0.384 | 0.375 | 0.375 | **0.542** | 0.486 | 0.306 | 0.500 | **0.542** |
| **Positive Sec/SPI** | ±0 | 0.766 | 0.807 | **0.820** | 0.809 | 0.745 | 0.671 | 0.752 | 0.803 | 0.814 | 0.816 | 0.785 | 0.732 | 0.813 |
| | ±1 | 0.779 | 0.818 | 0.830 | 0.826 | 0.750 | 0.679 | 0.763 | 0.819 | **0.838** | 0.838 | 0.796 | 0.741 | 0.832 |
| | ±2 | 0.794 | 0.833 | 0.848 | 0.841 | 0.769 | 0.692 | 0.772 | 0.834 | **0.855** | 0.853 | 0.816 | 0.756 | 0.852 |
| | ±3 | 0.812 | 0.839 | 0.854 | 0.859 | 0.777 | 0.698 | 0.778 | 0.840 | **0.861** | 0.859 | 0.825 | 0.774 | 0.860 |
| **Positive Sec/SPII** | ±0 | 0.870 | 0.901 | 0.911 | 0.890 | 0.800 | 0.872 | 0.866 | 0.912 | 0.881 | 0.898 | **0.929** | 0.903 | 0.921 |
| | ±1 | 0.870 | 0.901 | 0.911 | 0.890 | 0.802 | 0.872 | 0.866 | 0.912 | 0.881 | 0.898 | **0.929** | 0.903 | 0.921 |
| | ±2 | 0.871 | 0.901 | 0.911 | 0.890 | 0.802 | 0.872 | 0.866 | 0.912 | 0.881 | 0.898 | **0.929** | 0.903 | 0.921 |
| | ±3 | 0.871 | 0.901 | 0.911 | 0.890 | 0.802 | 0.872 | 0.866 | 0.912 | 0.881 | 0.898 | **0.929** | 0.903 | 0.921 |
| **Positive Sec/SPIII** | ±0 | **1.000** | **1.000** | 0.917 | **1.000** | 0.833 | **1.000** | 0.750 | **1.000** | **1.000** | **1.000** | 0.750 | 0.500 | 0.833 |
| | ±1 | **1.000** | **1.000** | 0.917 | **1.000** | 0.833 | **1.000** | 0.750 | **1.000** | **1.000** | **1.000** | 0.917 | 0.667 | 0.833 |
| | ±2 | **1.000** | **1.000** | 0.917 | **1.000** | 0.833 | **1.000** | 0.750 | **1.000** | **1.000** | **1.000** | 0.917 | 0.833 | 0.833 |
| | ±3 | **1.000** | **1.000** | 0.917 | **1.000** | 0.833 | **1.000** | 0.750 | **1.000** | **1.000** | **1.000** | 0.917 | 0.833 | **1.000** |
| **Positive Tat/SPI** | ±0 | 0.396 | 0.526 | 0.485 | 0.411 | 0.406 | 0.430 | 0.265 | 0.389 | 0.494 | 0.387 | 0.444 | 0.511 | **0.550** |
| | ±1 | 0.457 | 0.526 | 0.507 | 0.411 | 0.406 | 0.450 | 0.285 | 0.400 | 0.494 | 0.415 | 0.465 | 0.533 | **0.550** |
| | ±2 | 0.669 | 0.741 | 0.672 | 0.576 | 0.506 | 0.633 | 0.431 | 0.633 | 0.641 | 0.611 | 0.691 | 0.733 | **0.765** |
| | ±3 | 0.669 | 0.741 | 0.700 | 0.604 | 0.543 | 0.633 | 0.431 | 0.633 | 0.641 | 0.611 | 0.691 | 0.733 | **0.765** |
| **Positive Tat/SPII** | ±0 | **1.000** | 0.861 | 0.972 | 0.972 | 0.361 | 0.194 | 0.194 | 0.722 | 0.694 | **1.000** | 0.611 | 0.667 | 0.972 |
| | ±1 | **1.000** | 0.889 | 0.972 | 0.972 | 0.361 | 0.194 | 0.194 | 0.722 | 0.694 | **1.000** | 0.611 | 0.667 | 0.972 |
| | ±2 | **1.000** | 0.889 | 0.972 | 0.972 | 0.361 | 0.194 | 0.194 | 0.722 | 0.694 | **1.000** | 0.611 | 0.667 | 0.972 |
| | ±3 | **1.000** | 0.889 | 0.972 | 0.972 | 0.361 | 0.194 | 0.194 | 0.722 | 0.694 | **1.000** | 0.611 | 0.667 | 0.972 |
| **mean** | | 0.761 | 0.768 | 0.785 | 0.792 | 0.599 | 0.679 | 0.630 | 0.755 | **0.799** | 0.767 | 0.729 | 0.716 | 0.775 |
